# Gut IgA Enhances Systemic IgG Responses to Pneumococcal Vaccines Through the Commensal Microbiota

**DOI:** 10.1101/2021.04.29.439534

**Authors:** Cindy Gutzeit, Emilie K. Grasset, Dean B. Matthews, Paul J. Maglione, Giuliana Magri, Graham J. Britton, Lewis Tomalin, Marc Pybus, Sonia Tejedor Vaquero, Pavan K. Veeramreddy, Roser Tachó-Piñot, Andrea Martín Nalda, Marina García Prat, Monica Martinez Gallo, Romina Dieli-Crimi, José C. Clemente, Saurabh Mehandru, Mayte Suarez-Farina, Jeremiah J. Faith, Charlotte Cunningham-Rundles, Andrea Cerutti

## Abstract

The gut microbiota enhances systemic immunoglobulin G (IgG) responses to vaccines. However, it is unknown whether this effect involves IgA, a mucosal antibody that coats intestinal microbes. Here we found that gut IgA increased peripheral IgG responses to pneumococcal vaccines, as these responses were profoundly impaired in mice with global or mucosa-restricted IgA deficiency. The positive effect of IgA on vaccine-induced IgG production implicated gut bacteria. Indeed, IgG responses to pneumococcal vaccines were also defective in ex-germ free mice recolonized with gut microbes from mouse or human IgA-deficient donors. IgA exerted this IgG-enhancing effect by constraining the systemic translocation of intestinal commensal antigens, which caused chronic immune activation, including T cell overexpression of programmed death-1. This immune inhibitory receptor hindered vaccine-specific IgG production by eliciting functional B cell unresponsiveness, which was reverted by anti-programmed death-1 treatment. Thus, gut IgA is functionally interconnected with systemic IgG via intestinal microbes.

## INTRODUCTION

The intestinal mucosa is inhabited commensal bacteria, fungi, viruses and other microbial and eukaryotic species collectively referred to as gut microbiota, which modulate gut immunity, host metabolism and neuronal signaling (Belkaid and Hand, 2014; Sherwin et al., 2019; Sonnenburg and Backhed, 2016). The gut microbiota also modulates inflammatory, autoimmune and allergic disorders (Kamada et al., 2013; Renz and Skevaki, 2021; Ruff et al., 2020) as well as the outcome of immune interventions such as vaccination and cancer therapy (de Jong et al., 2020; Finlay et al., 2020), including treatment with checkpoint inhibitors of the programmed death-1 (PD-1) receptor (Gopalakrishnan et al., 2018). While these studies have mostly focused on the microbial component of the gut microbiota, this “prokaryotic organ” of our body further includes immunoglobulin A (IgA), a microbiota-binding antibody produced by gut B cells (Chen et al., 2020). Yet, the role of gut IgA in systemic immunity remains unknown.

IgA mostly derives from intestinal B cell responses to commensal bacteria, which function as major IgA inducers (Chen et al., 2020). These responses occur under homeostatic conditions through complementary T cell-dependent (TD) and T cell-independent (TI) pathways that yield distinct pools of IgAs (Bunker et al., 2015; Grasset et al., 2020). Conversely, IgA interaction with gut commensals shapes the topography, composition and immunometabolic properties of the microbiota (Macpherson et al., 2018). However, the impact of intestinal IgA on systemic IgG responses remains elusive.

Growing evidence indicates that IgA cooperates with IgG to enhance protection against both commensals and pathogens (Fadlallah et al., 2019; Koch et al., 2016; Rollenske et al., 2018). Accordingly, some patients with IgA deficiency (IGAD) develop mucosal infections in addition to gut dysbiosis (Catanzaro et al., 2019; Cunningham-Rundles and Ponda, 2005; Fadlallah et al., 2018). Of note, infections by common lung pathogens such as *Streptococcus pneumoniae* are usually controlled by specific IgG (Chen et al., 2020), which can be readily elicited by pneumococcal vaccines (Croucher et al., 2018). Remarkably, some IGAD patients with heterogeneous phenotypic traits show impaired systemic IgG responses to pneumococcal vaccines (Edwards et al., 2004; Lane and MacLennan, 1986). In addition, some IGAD patients exhibit a concomitant IgG subclass deficiency, whereas others develop a global and profound IgG depletion as they progress to common variable immunodeficiency (Aghamohammadi et al., 2008). Altogether, these observations suggest that IgA and namely gut IgA may be more functionally connected to IgG than commonly thought.

In agreement with this possibility, mouse studies have shown that the IgA-targeted motility protein flagellin from gut microbes enhances IgG responses to an influenza vaccine in addition to promoting IgA production (Cullender et al., 2013; Oh et al., 2014). Similarly, gut microbial metabolites amplify IgG in addition to IgA responses (Kim et al., 2016). Furthermore, mucosal immune adaptations to the microbiota, including IgA production, limit the extensive and potentially harmful host tissue response caused by unrestrained systemic penetration of and tissue exposure to gut metabolites (Uchimura et al., 2018). Notwithstanding this evidence, whether commensal-reactive intestinal IgA influences systemic IgG responses remains unknown.

Here we found that IgA increased IgG responses to both unconjugated and protein-conjugated pneumococcal vaccines through the gut microbiota. IgG responses were impaired in mice with global or mucosa-restricted IgA deficiency and in ex-germ free (GF) mice recolonized with gut microbes from IgA- deficient mice or IGAD patients. Gut IgA enhanced vaccine-specific IgG production by constraining the systemic translocation of commensal antigens. Indeed, the lack of IgA abnormally increased IgG responses to gut microbial products along with PD-1 expression, two processes linked to enhanced B and T cell activation, respectively. By interfering with IgG production in B cells, PD-1 inhibited newly induced IgG responses to pneumococcal vaccines, but this inhibition was rescued by treatment with anti-PD-1. IgA also sustained the abundance of gut microbiota-regulated systemic branched chain amino acids (BCAAs) (Pedersen et al., 2016), which amplified IgG production in B cells stimulated via TD or TI signaling programs *in vitro*. By showing that intestinal microbes functionally link gut IgA to systemic IgG, our findings unveil a novel facet of IgA biology and prompts to include IgA in studies assessing the impact of the intestinal microbiota on biological processes such as the response to vaccines.

## RESULTS

### IgA Enhances Systemic IgG Responses to TI Immunizations

The human vaccine Pneumovax23 encompasses unconjugated pneumococcal polysaccharides (PPS), generically referred to as capsular polysaccharides (CPS), from 23 highly pathogenic serotypes of *Streptococcus pneumonia* (Croucher et al., 2018). Similar to other microbial carbohydrates, PPS induce IgG by activating splenic marginal zone (MZ) B cells in a TI manner, as opposed to microbial proteins that activate follicular (FO) B cells in a TD manner (Cerutti et al., 2013; Croucher et al., 2018).

First, we determined systemic IgG responses to Pneumovax23 in IgA-deficient C57BL/6 *Igha*^−/−^ mice. These mice were generated by deleting 1) the I*α* exon, which initiates the germline transcription of both switch *α* region and constant *α* gene, the intronic switch *α* region, which guides class switching from IgM to IgA, and the 5’ half of the constant *α* gene, which encodes the constant region of IgA (Harriman et al., 1999). Wild type (WT) and *Igha*^−/−^ breeders were set up from heterozygous *Igha*^+/−^ parents to control for microbiota and genetic background variability between strains. WT used came from these breeders, unless indicated otherwise. Enzyme-linked immunosorbent assay (ELISA) showed that, compared to WT controls, *Igha*^−/−^ mice were unable to induce serum IgG3 to PPS at 3 and 7 days following intravenous (i.v.) immunization with Pneumovax23 (**Figure 1A**). Together with IgG3, IgM is a major component of humoral immunity to TI antigens in mice (Cerutti et al., 2013). *Igha*^−/−^ mice had increased PPS-specific IgM both under steady-state conditions and following Pneumovax23 immunization compared to WT controls (**Figure 1A**). Similar to Pneumovax23, mitomycin C-inactivated *Streptococcus pneumoniae* induced less bacteria-specific IgG3 but more bacteria-specific IgM in *Igha*^−/−^ mice compared to WT controls at day 7 following i.v. immunization (**Figure S1A**).

**Figure 1.**
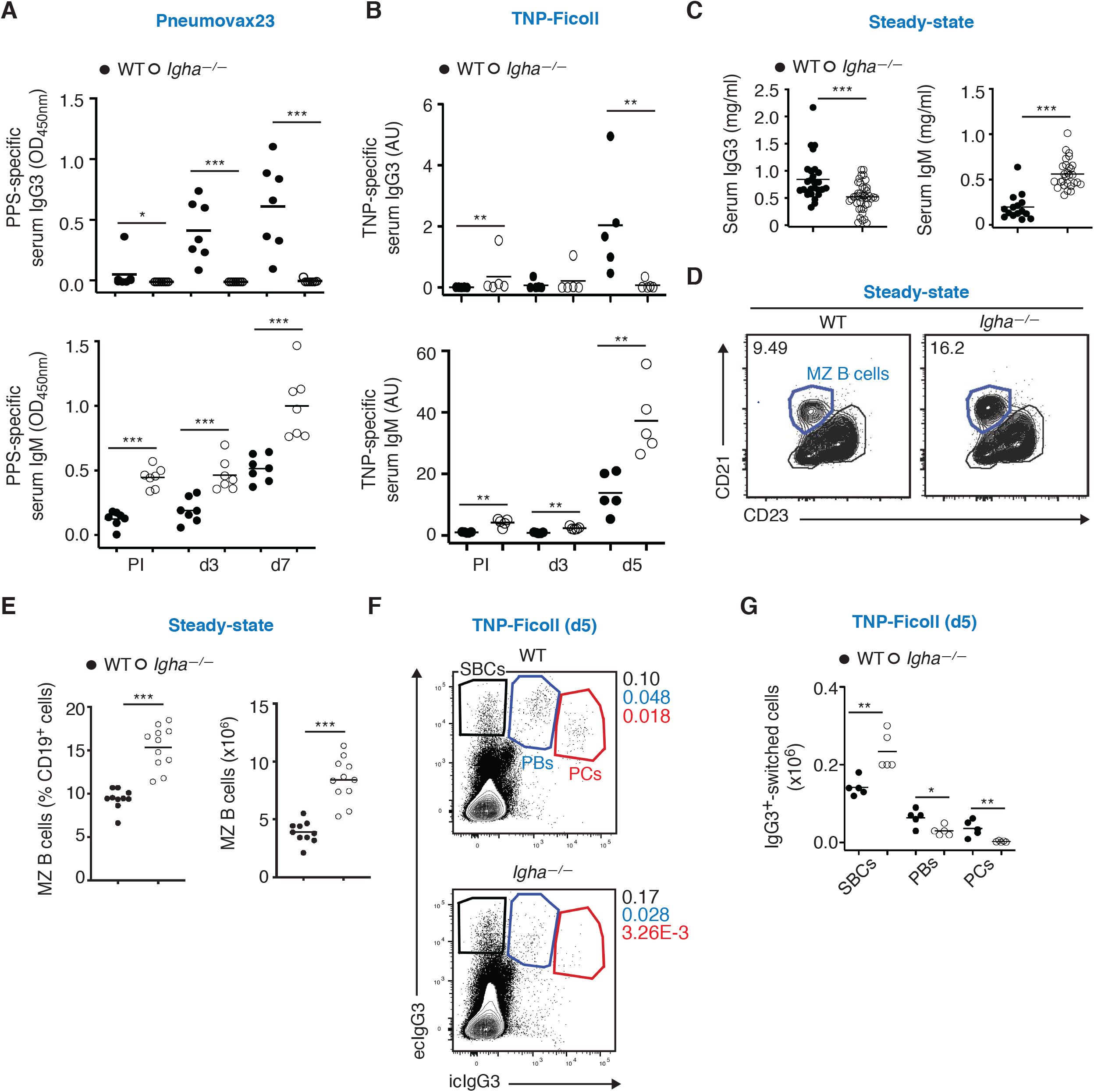
**IgA Enhances Systemic IgG Responses to TI Immunizations** (A) ELISA of serum IgG3 (top) and IgM (bottom) to PPS from Pneumovax23 in 7 WT (solid circles) or *Igha*^−/−^ (open circles) mice prior to immunization (PI) and 3 or 7 days following i.v. immunization with Pneumovax23. (B) ELISA of serum IgG3 (top) and IgM (bottom) to TNP from 5 WT or *Igha*^−/−^ mice before and 3 or 5 days following i.v. immunization with TNP-Ficoll. This experiment involved WT mice purchased from Jackson Laboratories. (C) ELISA of total serum IgG3 and IgM in 15-27 WT or 28-41 *Igha*^−/−^ mice at steady state. (D) Flow cytometry of CD21 and CD23 on splenic MZ B cells from representative WT or *Igha*^−/−^ mice. Numbers indicate MZ B cell frequency (%) within the CD19^+^ population. (E) Frequency (% of CD19^+^ cells; left) and absolute number (right) of splenic CD21^high^CD23^low^ MZ B cells from 10 WT mice or 11 *Igha*^−/−^ mice (E). (F, G) Flow cytometry of intracellular (ic) IgG3 and extracellular (ec) IgG3 from splenic ecIgG3^+^icIgG3^−^ switched B cells (SBCs), ecIgG3^+^icIgG3^+^ plasmablasts (PBs) and ecIgG3^lo^icIgG3^+^ PCs of representative WT or *Igha*^−/−^ mice (F) as well as absolute number (G) of SBCs, PBs and PCs from 5 WT or 5 *Igha*^−/−^ mice 5 days following i.v. immunization with TNP-Ficoll. Numbers in (F) indicate frequency within total cells. These experiments involved WT mice purchased from Jackson Laboratories. Data show representative flow plots (D, F) or summarize results from one (A, B, E, G) or 5-6 experiments (C). Results are shown with mean; a two-tailed unpaired Student’s t-test was performed when data followed a Gaussian distribution, otherwise a Mann-Whitney test was used to determine significance. *p < 0.05, **p < 0.01, ***p <0.001. See also Figure S1.

Next, we ascertained whether IgA deficiency impaired IgG3 responses to TI antigens different from unconjugated PPS. Compared to WT controls, *Igha*^−/−^ mice showed no 2,4,6-trinitrophenyl (TNP)- specific IgG3 but increased TNP-specific IgM at steady state and 3 or 5 days following i.v. immunization with TNP-Ficoll (**Figure 1B**), a haptenated polysaccharide structurally mimicking native CPS. While TNP-Ficoll activates MZ and B-1 B cells by crosslinking the B cell antigen receptor (BCR), TNP- lipopolysaccharide (LPS) activates MZ and B-1 B cells by co-engaging Toll-like receptor 4 (TLR4) and BCR (Pone et al., 2012). Compared to WT controls, *Igha*^−/−^ mice induced less TNP-specific IgG3 but more TNP-specific IgM 3 and 7 days following i.v. immunization with TNP-LPS (**Figure S1B**).

We then wondered whether IgA deficiency globally impaired IgG3 production by B cells, including MZ B cells. At steady state, total serum IgG3 was readily detectable in *Igha*^−/−^ mice, although its concentration was reduced compared to WT controls (**Figure 1C**). This finding ruled out a global impairment of IgM-to-IgG3 class switching and IgG3 secretion in B cells from *Igha*^−/−^ mice. As reported by an earlier study (Harriman et al., 1999), these mice also showed more total serum IgM at steady state compared to WT controls (**Figure 1C**), which implied that IgA deficiency was not associated with a general impairment of B cell activation and antibody production.

We next wondered whether IgA deficiency caused a depletion of MZ B cells and/or impaired the capture of TI antigens by MZ B cells and neighboring macrophages (Cerutti et al., 2013). Flow cytometry showed an increase in splenic MZ B cells in *Igha*^−/−^ mice compared to WT controls at steady state (**Figure 1D** and **1E**). In addition, tissue immunofluorescence analysis detected comparable capture of TNP-Ficoll by MZ-based B cells and metallophilic macrophages from *Igha*^−/−^ mice and WT controls at 30 minutes following i.v. immunization along with comparable shuttling into follicles at 3 hours following immunization (**Figure S1C**). Altogether, these data suggest that the impaired response to TI antigens in *Igha*^−/−^ mice does not stem from gross perturbations of MZ B cells.

Finally, we determined whether IgA deficiency impaired IgG3 class switching and plasma cell (PC) differentiation in splenic B cells exposed to TI antigens. Compared to WT controls, splenic IgG3 class-switched B cells actually increased 5 days following i.v. immunization of *Igha*^−/−^ mice with TNP-Ficoll, whereas IgG3 class-switched plasmablasts and PCs decreased (**Figure 1F** and **1G**). This decrease was not due to an intrinsic B cell defect, as splenic B cells from WT and *Igha*^−/−^ mice cultured with the TI antigen LPS for 4 days comparably differentiated to plasmablasts (**Figure S1D** and **S1E**). Thus, IgA enhances systemic IgG responses to TI immunogens and may do so through a B cell-extrinsic mechanism involving increased PC differentiation and/or survival.

### IgA Increases Systemic IgG Responses to TD Immunizations

Next, we determined whether IgA deficiency impaired systemic IgG responses to the TD conjugated pneumococcal vaccine Prevnar13, which encompasses protein-conjugated PPS from 13 highly pathogenic serotypes of *Streptococcus pneumonia* (Croucher et al., 2018). Compared to WT controls, *Igha*^−/−^ mice induced less IgG1 to PPS from the Prevnar13 vaccine 7 and 28 days following i.v. immunization (**Figure 2A**). However, this decrease was not associated with gross quantitative alterations of splenic class-switched B cell, plasmablasts and PCs (**Figure 2B** and **Figure S2A**). We then verified whether IgA deficiency impaired the magnitude and affinity of systemic IgG1 responses to 4-hydroxy-3-nitrophenyl- ovalbumin (NP-OVA), a commonly used TD immunogen. Compared to WT controls, *Igha*^−/−^ mice induced less IgG1 with either high (NP7) or low (NP23) affinity 7, 14, 21, 28 or 62 days following i.p. immunization with NP-OVA and alum (**Figure 2C** and **Figure S2B**). In immunized *Igha*^−/−^ mice, the reduction of IgG1 affinity maturation was most evident at day 7 and 14 and remained significant up to day 28 (**Figure 2D**).

**Figure 2.**
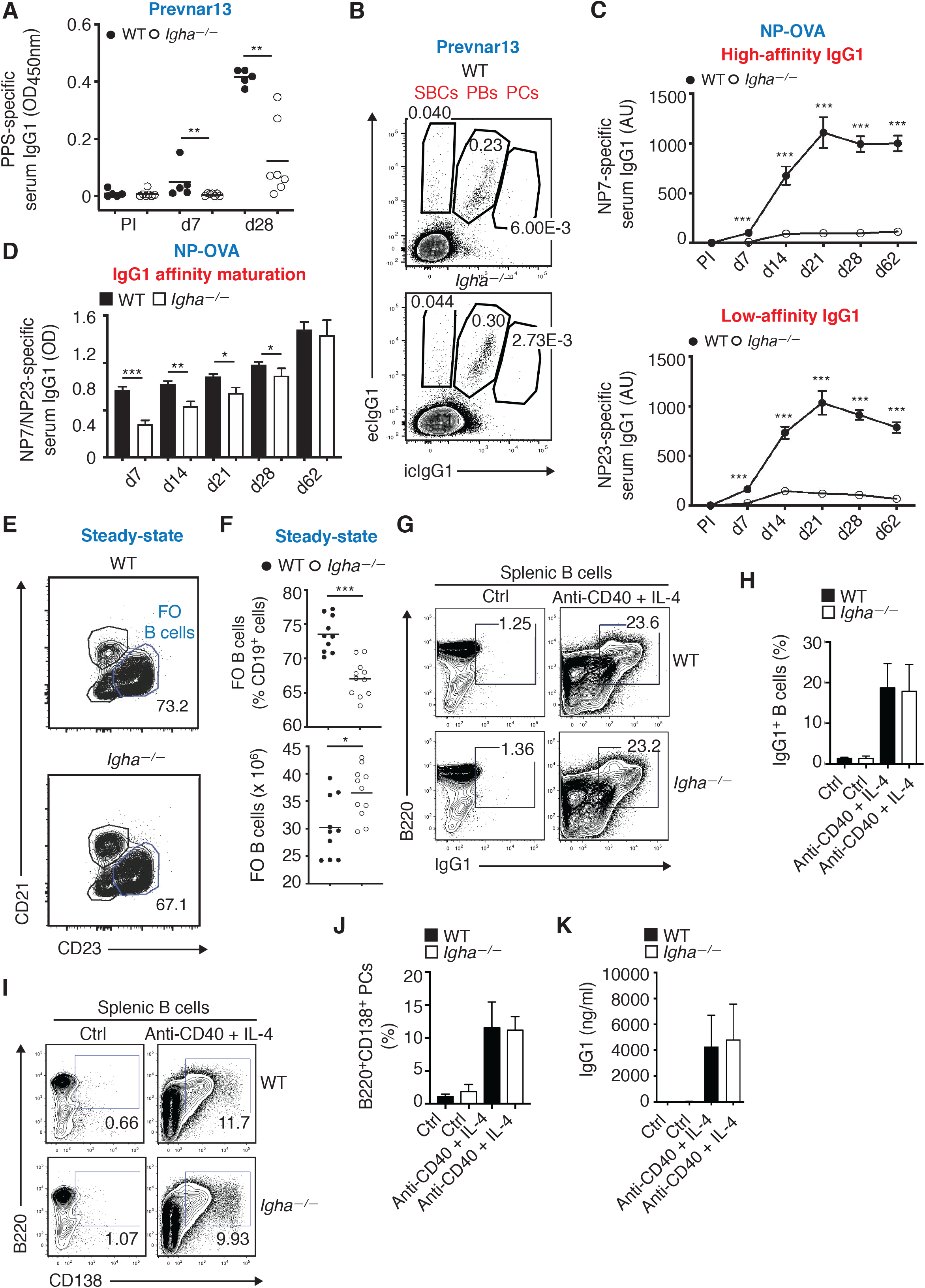
**IgA Increases Systemic IgG Responses to TD immunizations** (A) ELISA of serum IgG1 to PPS from 5 WT or 7 *Igha*^−/−^ mice prior to immunization (PI) and 7 or 28 days following i.v. immunization with Prevnar13. (B) Flow cytometry of intracellular (ic) IgG1 and extracellular (ec) IgG1 from splenic ecIgG1^+^icIgG1^−^ switched B cells (SBCs), ecIgG1^+^icIgG1^+^ plasmablasts (PBs) and ecIgG1^lo^icIgG1^+^ PCs of representative WT or *Igha*^−/−^ mice 28 days following i.v. immunization with Prevnar13. Numbers indicate frequency (% of B220^+^ cells) of SBCs, PBs and PCs. (C) ELISA of serum high-affinity (NP7-BSA) IgG1 and low-affinity (NP23-BSA) IgG1 from 13 WT or 11 *Igha*^−/−^ mice before immunization (PI) and 7, 14, 21, 28 or 62 days following i.p. immunization with NP15-OVA and alum. (D) Affinity maturation of IgG1 calculated as a ratio of IgG1 specific to NP7-BSA versus IgG1 specific to NP23-BSA in 13 WT or 11 *Igha^-/-^* mice 7, 14, 21, 28 or 62 days following i.p. immunization with NP15- OVA and alum. (E, F) Flow cytometry of CD21 and CD23 on splenic FO B cells from representative WT or *Igha*^−/−^ mice at steady state (E) as well as frequency (% of CD19+ cells) and absolute number of CD21^low^CD23^high^ FO B cells from 10 WT or 11 *Igha*^−/−^ mice (F). (G, H) Flow cytometry of B220 and IgG1 on purified splenic B cells from representative WT or *Igha*^−/−^ mice incubated with medium alone (ctrl) or anti-CD40 and IL-4 for 6 days (G) as well as frequency (% of live cells) of splenic IgG1^+^B220^+^ B cells from 6 WT or *Igha*^−/−^ mice cultured as above (H). (I, J) Flow cytometry of B220 and CD138 on purified splenic B cells from representative WT or *Igha*^−/−^ mice incubated with medium alone (ctrl) or anti-CD40 and IL-4 for 6 days (I) as well as frequency (% of live cells) of splenic B220^+^CD138^+^ PBs from 6 WT or *Igha*^−/−^ mice cultured as above (J). (K) ELISA of IgG1 secreted by purified splenic B cells from representative 4 WT or *Igha*^−/−^ mice incubated with medium alone (ctrl) or anti-CD40 and IL-4 for 6 days. Data show one representative result (B, E, G, I) or summarize results from either a single (F), two (A, C, D, K), or three experiments (H, J) Results are shown with mean (A, F) or mean ± s.e.m. (C, D, H, J, K); a two-tailed unpaired Student’s t-test was performed when data followed a Gaussian distribution, otherwise a Mann-Whitney test was used to determine significance. *p < 0.05, **p < 0.01, ***p <0.001. See also Figure S2.

We also wondered whether IgA deficiency depleted FO B cell precursors of IgG1-secreting cells and found that the absolute number but not the frequency of these cells was increased in steady-state *Igha*^−/−^ mice compared to WT controls (**Figure 2E** and **2F**). We further evaluated whether IgA deficiency caused B cell-intrinsic alterations of TD-induced IgG1 class switching and IgG1 secretion. However, we found that splenic B cells from *Igha*^−/−^ mice induced surface IgG1, differentiated into PCs, and secreted IgG1 as much as splenic B cells from WT controls upon incubation for 6 days with T cell-associated stimuli such as an agonistic antibody to CD40 mimicking CD40 ligand (CD40L) and interleukin-4 (IL-4) (**Figure 2G**- **K**). Finally, we determined whether IgA deficiency altered splenic T cells and dendritic cells (DCs), which are essential to initiate TD antibody responses (Crotty, 2019). Compared to WT controls, *Igha*^−/−^ mice showed overlapping frequencies and absolute numbers of total T cells as well as CD4^+^, CD8^+^ or CD4^−^CD8^−^ T cell subsets, except that the frequency of CD4^−^CD8^−^ T cells was slightly decreased and the absolute number of CD4^+^ T cells was slightly increased (**Figure S2C**). Compared to WT controls, *Igha*^−/−^ mice also showed overlapping frequencies and absolute numbers of total DCs as well as CD4^+^CD8^−^, CD4^−^CD8^+^ or CD4^−^CD8^−^ DC subsets (**Figure S2D**). Thus, IgA may enhance systemic IgG responses to TD immunogens through a B cell-extrinsic mechanism operating upstream of PC differentiation.

### IgA Helps Systemic IgG Responses to Vaccines Through Gut Commensals

Next, we asked whether the selective lack of intraluminal secretory IgA (SIgA) from mucosal membranes was sufficient to phenocopy global IgA deficiency. SIgA, including gut SIgA, is absent in *Pigr*^−/−^ mice, which lack the polymeric Ig receptor (pIgR) that transports dimeric IgA across mucosal epithelial cells to generate SIgA (Chen et al., 2020; Johansen et al., 1999). Similar to *Igha*^−/−^ mice, *Pigr*^−/−^ mice induced less serum IgG1 to PPS from the Prevnar13 vaccine 14 and 21 days following i.p. immunization compared to WT controls (**Figure 3A**).

**Figure 3.**
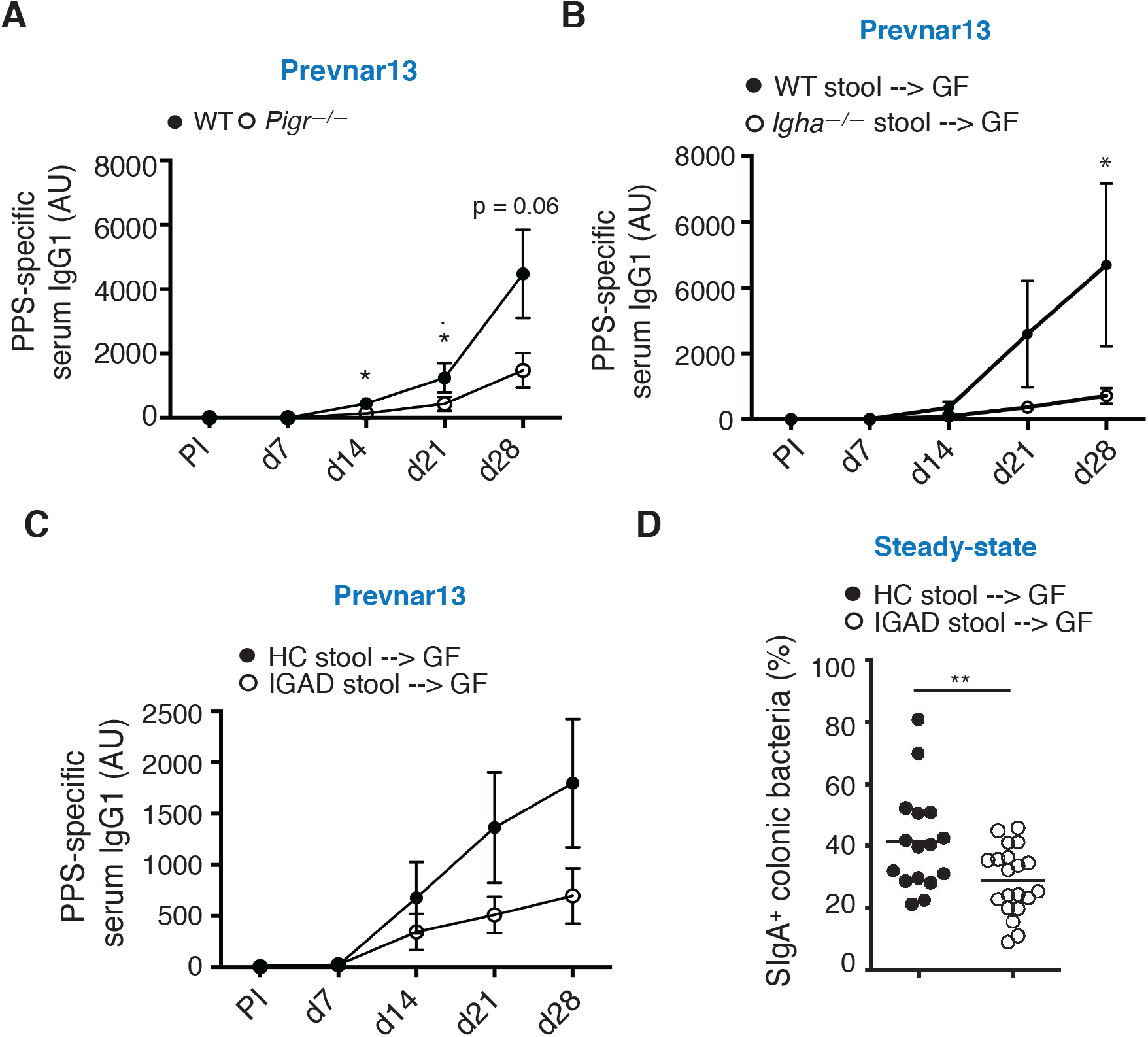
**IgA Helps Systemic IgG Responses to Vaccines through Gut Commensals** (A) ELISA of serum IgG1 to PPS from 8 WT mice or *Pigr*^−/−^ mice prior to immunization (PI) and 7, 14, 21 or 28 days following i.p. immunization with Prevnar13. (B) ELISA of serum IgG1 to PPS PI or 7, 14, 21 or 28 days following i.p. immunization with Prevnar13 of ex-GF mice obtained by reconstituting 6-7 GF recipient mice with cecal content from either an SPF WT or an SPF *Igha*^−/−^ donor mouse. Recipient mice were vaccinated two weeks following reconstitution. (C) ELISA of serum IgG1 to PPS prior to immunization or 7, 14, 21 or 28 days following i.p. immunization with Prevnar13 of ex-GF mice obtained by reconstituting 12 or 18 GF recipient mice with fecal material from 4 human HCs or 4 IGAD patients, respectively. Immunization was performed four weeks following reconstitution of the gut microbiota. (D) Flow cytometry of IgA^+^ gut bacteria from 16 or 20 ex-GF mice reconstituted with fecal material from 4 healthy controls (HC) and 4 IGAD donors, respectively. Measurements were performed four weeks following reconstitution of the gut microbiota. Results are from 1 (B) or 2 (A, C, D) independent experiments. Data are shown with mean (D) or mean ± s.e.m (A, B, C); a two-tailed unpaired Student’s t-test was performed when data followed a Gaussian distribution, otherwise a Mann-Whitney test was used. *p < 0.05, **p < 0.01. See also Figure S3.

Having shown that mucosa-restricted IgA deficiency was sufficient to hamper systemic IgG1 responses to Prevnar13 and knowing that SIgA targets gut commensals, we determined whether intestinal microbes from IgA-sufficient or IgA-deficient mice differentially induced systemic IgG1 responses to Prevnar13. Compared to ex-GF mice obtained by reconstituting GF recipients with fecal bacteria from WT donor mice, ex-GF mice obtained by reconstituting GF recipients with fecal bacteria from *Igha*^−/−^ donors induced less IgG1 to PPS 28 days following i.p. immunization with Prevnar13 (**Figure 3B**).

We then evaluated whether human gut commensals from healthy controls or IGAD patients enhanced systemic IgG1 responses to Prevnar13. This experiment was performed by taking advantage of fecal samples from a New York City (NYC) cohort of adult IGAD patients (**Table S1**). Compared to control ex-GF mice colonized with fecal bacteria from healthy human donors, ex-GF mice colonized with fecal bacteria from IGAD patients showed a clear trend toward decreased serum IgG1 to PPS 21 or 28 days following i.p. immunization with Prevnar13, albeit this reduction failed to reach statistical significance (**Figure 3C**). Then, we hypothesized that human IgA could enhance gut IgA responses to the local microbiota in addition to systemic IgG responses to vaccines. Compared to control ex-GF mice colonized with fecal bacteria from healthy human donors, ex-GF mice colonized with fecal bacteria from adult IGAD patients showed decreased SIgA-coated gut commensals (**Figure 3D**). This finding indicates that, compared to gut bacteria from controls, gut microbes from IGAD patients induce SIgA less efficiently.

Finally, we tested whether lateral transmission of IgA-coated gut commensals restored IgG1 responses to pneumococcal vaccines. Compared to control *Igha*^+/+^ littermates, co-housed *Igha*^−/−^ mice retained decreased serum IgG1 to PPS from Pneumovax23 or Prevnar13 vaccines 28 days following i.p. immunization (**Figure S3A** and **S3B**). Thus, IgA may enhance systemic IgG responses to pneumococcal vaccines and gut IgA responses to commensals through a mechanism involving the gut microbiota.

### IgA Restrains Systemic IgG Responses to Translocated Gut Antigens

To further elucidate how IgA augments systemic IgG1 responses to vaccines, we first ascertained the effect of IgA on global IgG1 production under steady-state conditions. Consistent with an earlier study (Harriman et al., 1999), we found that total serum IgG1 was increased in *Igha*^−/−^ mice compared to WT controls (**Figure 4A**). Therefore, the impaired IgG1 response to Prevnar13 or NP-OVA was not due to an overall decrease of IgG1 production. Given the strong dependence of IgG1 responses on B cell activation by IL-4 (Shan et al., 2018), this finding raised the possibility that IgA deficiency also enhanced systemic IL-4 production. Accordingly and similar to an earlier study (Harriman et al., 1999), we found that serum from steady-state *Igha*^−/−^ mice had more IL-4 compared to WT controls (**Figure 4B**).

**Figure 4.**
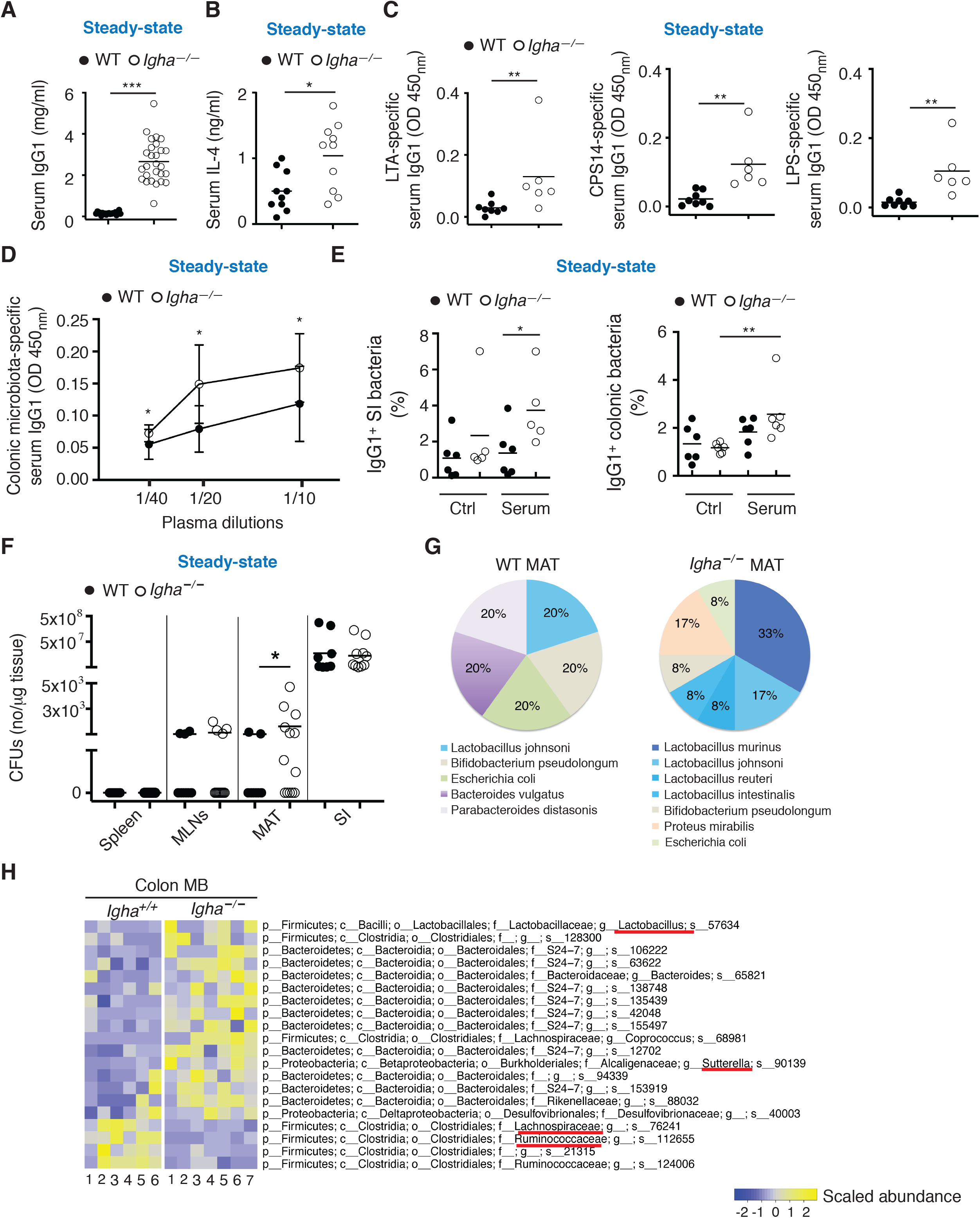
**IgA Restrains Systemic IgG Responses to Translocated Gut Antigens** (A) ELISA of total serum IgG1 from 20 WT or 27 *Igha*^−/−^ mice at steady state. (B) ELISA of serum IL-4 from 10 WT or 10 *Igha*^−/−^ mice at steady state. (C) ELISA of serum IgG1 to lipoteichoic acid (LTA) from *Staphylococcus aureus*, CPS14 from *Streptococcus pneumoniae* or LPS from *Salmonella typhimurium* in 8 WT or 6 *Igha*^−/−^ littermate mice at steady state. (D) ELISA of serum IgG1 reactive to paired colonic microbial antigens from 15 WT or 16 *Igha*^−/−^ mice at steady state (fecal and serum samples from same mouse). (E) Flow cytometry of IgG1-coated small intestinal (SI) (left) or colonic (right) bacteria from 6 WT or 5-6 *Igha*^−/−^ mice with or without (ctrl) incubation with paired serum. (F) Quantification of CFUs of anaerobic bacteria from homogenates of spleen, MLNs, mesenteric adipose tissue (MAT) and SI from 8-13 WT or 10-13 *Igha*^−/−^ mice at steady state. (G) Taxonomic classification of anaerobic bacterial colonies isolated in the MAT from 2 WT or 7 *Igha*^−/−^ mice, as determined by mass spectrometry. (H) 16S rRNA gene sequencing analysis of microbiota (MB) from colon feces of 6 WT or 7 *Igha*^−/−^ co-housed littermate mice at steady state. Bacterial families detected in (G) or potentially relevant to gut homeostasis, including persistence of intraluminal SIgA, are underlined in red. Results summarize 2-3 (A, E) or 5 (F, G) independent experiments and one experiment (B, C, D, H). Data are presented as mean ± s.e.m.; a two-tailed unpaired Student’s t-test was performed when data followed a Gaussian distribution, otherwise a Mann-Whitney test was used. *p < 0.05, **p < 0.01, ***p <0.001. See also Figure S4.

Knowing that IgA limits the systemic penetration of gut antigens (Chen et al., 2020), we next ascertained whether IgA deficiency increased the systemic IgG1 reactivity to gut microbial products. Indeed, compared to WT controls, *Igha*^−/−^ mice at steady state showed more serum IgG1 to lipoteichoic acid (LTA) from *Staphylococcus aureus*, CPS14 from *Streptococcus pneumoniae*, LPS from *Salmonella typhimurium* as well as CPS9 from *Streptococcus pneumoniae* (**Figure 4C** and **Figure S4A**).

Moreover, we determined whether IgA deficiency increased peripheral IgG1 responses to gut microbial antigens. ELISA and flow cytometry showed that, compared to WT controls, *Igha*^−/−^ mice had increased serum IgG1 to gut bacteria, particularly bacteria from colon (**Figure 4D** and **4E**). Then, we verified whether IgA deficiency perturbs IgG1 responses also in the gut lumen. Similar to an earlier study (Harriman et al., 1999), *Igha*^−/−^ mice showed increased total IgG1 in addition to increased total IgM in feces from both small intestine and colon compared to WT controls (**Figure S4B** and **S4C**). Given that increased luminal IgG may originate from gut inflammation (Castro-Dopico et al., 2019), we also verified whether IgA deficiency caused intestinal inflammation. However, microscopic analysis showed neither small intestine nor colon inflammation in *Igha*^−/−^ mice (**Figure S4D**).

Having shown that IgA deficiency abnormally increased systemic IgG1 production to gut commensal antigens, we next evaluated whether this immune anomaly stemmed from augmented bacterial breaching of the intestinal barrier. Similar to mesenteric adipose tissue from patients with Crohn’s disease (Ha et al., 2020), mesenteric adipose tissue but not mesenteric lymph nodes (MLNs) or spleen tissue from *Igha*^−/−^ mice yielded more colony-forming units (CFUs) upon incubation on agar plates placed in an anaerobic chamber compared to WT controls (**Figure 4F** and **Figure S4E**). Mass spectrometry showed that, compared to WT controls, *Igha*^−/−^ mice displayed an increase of *Lactobacillus* (*L.*) species, including *L. murinus*, *L. reuteri* and *L. intestinalis* in the mesenteric adipose tissue (**Figure 4G** and **Figure S4F**).

Aside from confirming *Lactobacillus* expansion, a subsequent experiment involving 16S ribosomal RNA (rRNA) gene sequencing showed an increased abundance of the IgA-degrading bacterial genus *Sutterella* (Kaakoush, 2020) in the fecal microbiota from *Igha*^−/−^ mice compared to co-housed *Igha^+/+^* littermate controls (**Figure 4H**). Similar to IGAD patients (Fadlallah et al., 2018), *Igha*^−/−^ mice showed gut microbiota depletion of *Ruminococcacae* and *Lachnospiraceae*, which may be implicated in homeostatic IgA responses to commensals (Chen et al., 2020; Magri et al., 2017). Thus, IgA regulates the composition of the gut microbiota and attenuates the systemic translocation of gut microbial antigens, which otherwise cause unrestrained peripheral induction of commensal-reactive IgG.

### IgA Limits Systemic Expansion of GC B Cells

Considering the importance of T follicular helper (TFH) cells in B cell activation and IgG1 production (Crotty, 2019), we explored whether germinal centers (GCs) from gut-associated Peyer’s patches, MLNs and spleen of *Igha*^−/−^ mice showed any B and/or T cell anomalies. Compared to WT controls, steady-state *Igha*^−/−^ mice showed enlarged Peyer’s patches and MLNs but normally sized spleen upon macroscopic analysis (**Figure 5A** and **Figure S5A**). Accordingly, Peyer’s patches and MLNs from *Igha*^−/−^ mice had an increased cellularity, while the spleen did not (**Figure 5B**). However, flow cytometry detected increased GC B cells and TFH cells not only in Peyer’s patches and MLNs, but also in the spleen from *Igha*^−/−^ mice (**Figure 5C-5F**).

**Figure 5.**
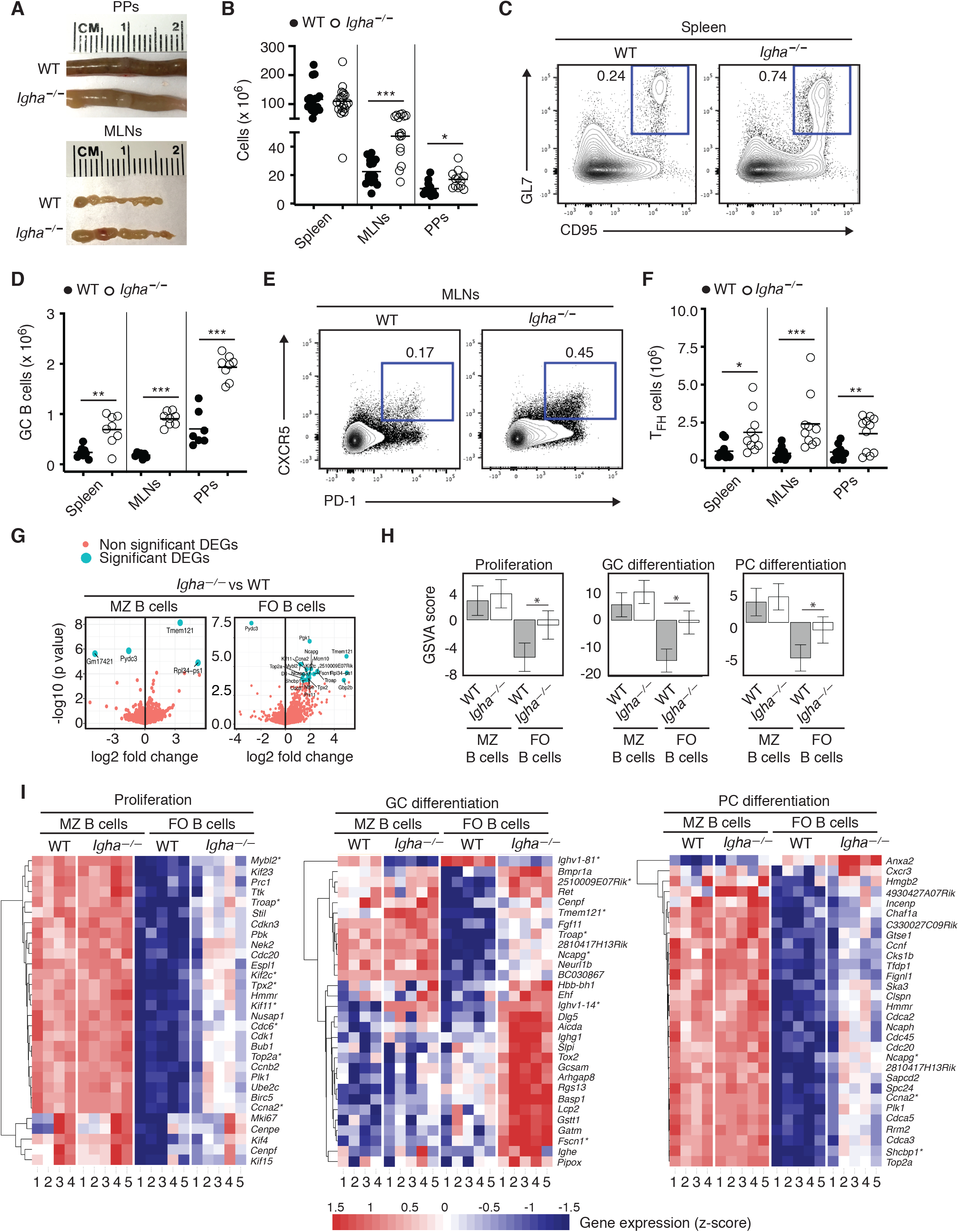
**IgA Limits Systemic Expansion of GC B Cells Along with PC Differentiation** (A) Images of Peyer’s patches and MLNs from representative WT or *Igha*^−/−^ mice at steady state. (B) Number of total cells in spleen, MLNs and Peyer’s patches from 11-19 WT or 12-20 *Igha*^−/−^ mice at steady state. (C, D) Flow cytometry analysis of splenic CD95^+^GL7^+^ GC B cells from representative WT or *Igha*^−/−^ mice at steady state (C) and absolute number of these cells in spleen, MLNs and Peyer’s patches from 7-8 WT or 8 *Igha*^−/−^ mice at steady state. GC B cells were analyzed from an initial CD45^+^B220^+^ gate. (E, F) Flow cytometry analysis of PD-1^+^CXCR5^+^ TFH cells from MLNs of representative WT or *Igha*^−/−^ mice at steady state (E) and absolute number of these TFH cells in spleen, MLNs and Peyer’s patches from 10-11 WT or 11 *Igha*^−/−^ mice (F). TFH cells were analyzed from an initial CD45^+^CD19^−^TCR*β*^+^CD4^+^ gate. (G) Volcano plot analysis of differentially expressed genes (DEGs) identified by RNA-seq in splenic MZ and FO B cell from 4-5 WT or 5 *Igha*^−/−^ mice at steady state. (H) Gene set variation analysis (GSVA) of proliferation (G2-M checkpoint), GC differentiation and PC differentiation gene signatures identified by RNA-seq and differentially expressed by splenic FO but not MZ B cells from 4 WT or 5 *Igha*^−/−^ mice at steady state. (I) Heat maps show z-score of gene-expression for top 30 differentially expressed genes from proliferation (left), GC differentiation (mid) and PC differentiation (right) gene sets identified by RNA-seq to be significantly differentially expressed (as in H) by splenic FO but not MZ B cells from 4-5 WT or 5 *Igha*^−/−^ mice at steady state. Asterisks indicate highly differentially expressed genes discussed in the text. The scale bar shows color coding (blue, white, red) for z-score. Data show representative images or flow plots (A, C, E), summarize 3 (D, F) or 6 (B) independent experiments, or one experiment (G, H, I). Data are presented with mean (D, F) or mean GSVA score and 95% confidence interval (H); a two-tailed unpaired Student’s t-test was performed when data were determined to follow a Gaussian distribution (B), otherwise a Mann-Whitney test was used (D, F), or limma modeling (H). *p < 0.05, **p < 0.01, ***p <0.001. See also Figure S5.

Given that gut IgA starts influencing the intestinal microbiota soon after birth and that gut bacteria stimulate early post-natal immune development (Gomez de Aguero et al., 2016; Planer et al., 2016), we assessed whether IgA deficiency caused an early onset of mucosal and systemic immune anomalies. Indeed, we found that both size and cellularity of MLNs as well as total systemic IgG1 were already increased in 5 week-old *Igha*^−/−^ mice compared to WT controls and that this increase became more pronounced in 16 week-old *Igha*^−/−^ mice (**Figure S5B-5D**). Unlike total IgG1, specific IgG3 to TNP was virtually absent in 5 week-old *Igha*^−/−^ mice 5 days following i.p. immunization with TNP-Ficoll (**Figure S5E**). Thus, IgA may enhance systemic IgG responses to immunogens by constraining the stimulation of mucosal and systemic B cells by gut microbial antigens since early post-natal life.

### IgA Controls Systemic FO B Cell Differentiation into PCs

Having shown that IgA deficiency increased splenic FO B cell numbers but impaired IgG responses to TD immunogens, we set up to characterize the global transcriptome of splenic FO cells from control WT and *Igha*^−/−^ mice with the aim of gaining new insights into how translocated gut antigens may impair IgG production to vaccines in the absence of IgA. Splenic MZ B cells were also included in this study, as IgA deficiency impaired IgG responses to TI immunogens as well.

As shown by volcano plot analysis of RNA-sequencing (RNA-seq) data, FO B cells from *Igha*^−/−^ mice showed more differentially expressed genes than MZ B cells did when compared to WT controls (**Figure 5G**). In addition, gene set variation analysis (GSVA) showed that, when compared to WT controls, FO B cells from *Igha*^−/−^ mice expressed significantly more gene sets linked to processes central to humoral immunity, including B cell proliferation, GC differentiation and PC differentiation (**Figure 5H**).

As shown by heat maps of RNA-sequencing (RNA-seq) data (**Figure 5I**), these increased gene sets included transcripts listed in the Hallmark_G2M_checkpoint gene set or described by a recent study (Shi et al., 2015) and implicated in B cell proliferation (e.g., *Mybl2*, *Troap*, *Kif2c*, *Tpx2*, *Kif11, Cdc6, Top2a, Ccna2*), GC differentiation (e.g., *Tmem121, 2510009E07Rik*, *Troap*, *Ncapg*, *Fscn1*) and PC differentiation (e.g., *Ncapg*, *Ccna2*, *Shcbp1*). Thus, IgA may enhance systemic IgG responses to vaccines by constraining the differentiation of systemic FO B cells into IgG-secreting PCs via the GC reaction. A corollary conclusion of these results is that IgA deficiency triggers B cell overstimulation by preferentially activating FO rather than MZ pathways.

### IgA Constrains Systemic T Cell Expression of the Immune Inhibitor PD-1 and Anti-PD-1 Rescues the Vaccine-Specific IgG Production in IgA Deficiency

Having shown that IgA constrains IgG responses to gut microbial antigens through a mechanism that starts early in life, we hypothesized that, in the absence of IgA, the abnormal systemic translocation of these endogenous antigens could impair IgG responses to vaccines by chronically activating B cells, potentially causing anergy. This B cell state refers to functional unresponsiveness caused by early and persistent exposure to self-antigens (Sabouri et al., 2016). Accordingly, GSVA indicated that, compared to WT controls, splenic FO but not MZ B cells from *Igha*^−/−^ mice were enriched in genes linked to B cell anergy (**Figure 6A**). As shown by a heat map of RNAseq data, this gene set, selected based on recently published findings (Sabouri et al., 2016), included *Cenpv*, *Lef1*, *Nefh*, *Fabp5* and *Tgif1* (**Figure 6B**).

**Figure 6.**
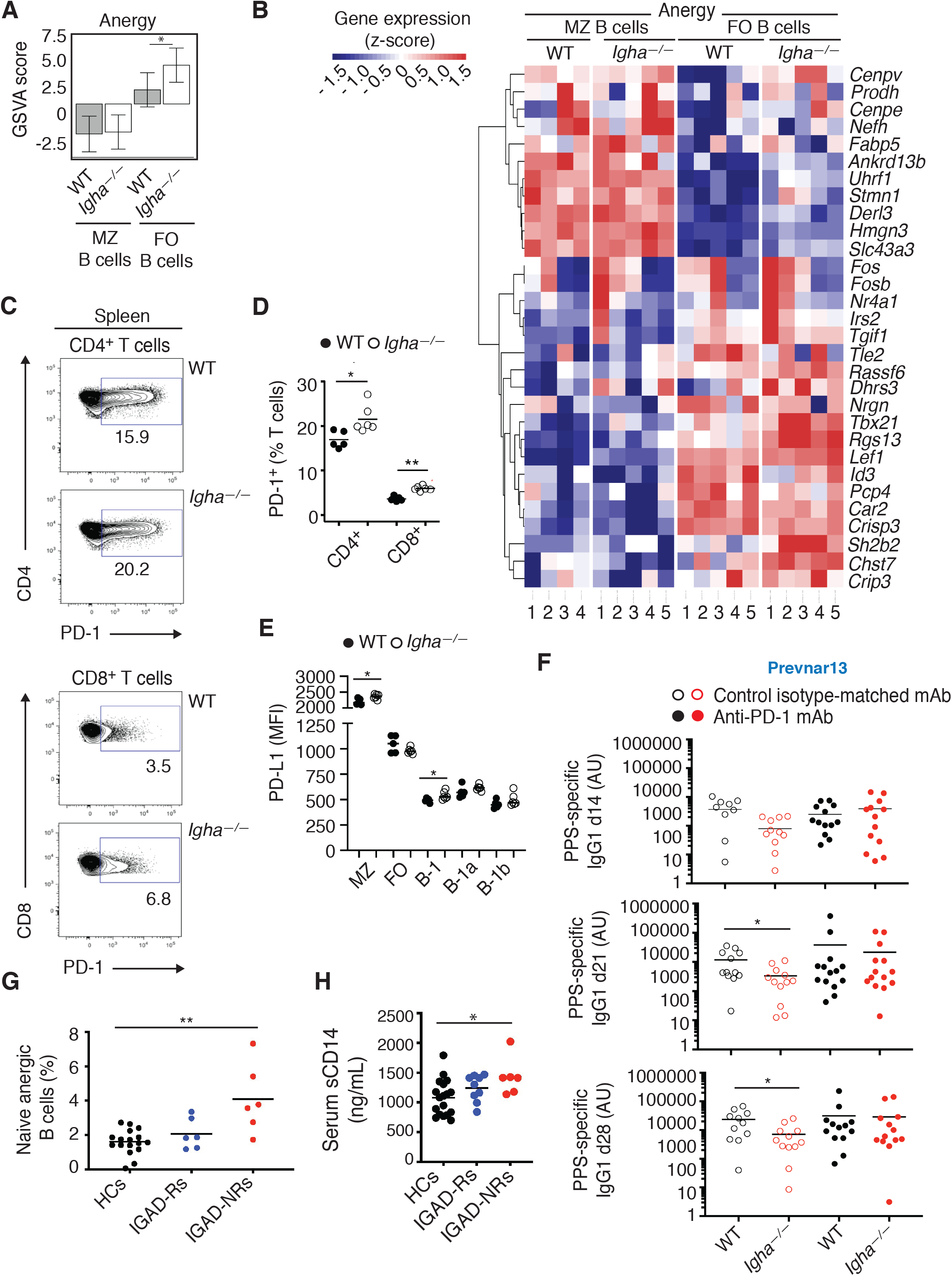
**IgA Constrains Systemic T Cell Expression of the immune inhibitor PD-1 and Anti-PD-1 Rescues Vaccine-Specific IgG Production in IgA Deficiency** (A) Gene set variation analysis (GSVA) of anergy gene signatures identified by RNA-seq and differentially expressed by splenic MZ and FO B cells from 4-5 WT or 5 *Igha*^−/−^ mice at steady state. (B) Heat map shows z-score of gene-expression for top 30 differentially expressed anergy genes identified by RNA-seq from gene set differentially expressed (in A) by splenic FO but not MZ B cells from 4-5 WT or 5 *Igha*^−/−^ mice at steady state. Asterisks indicate highly differentially expressed genes discussed in the text. The scale bar shows color coding (blue, white, red) for z-score. (C) Flow cytometry of PD-1 and CD4 (top) or CD8 (bottom) on splenic CD3^+^ T cells from representative WT or *Igha*^−/−^ mice at steady state. Numbers indicate frequency (%) of PD-1-expresing cells. (D) Frequency (%) of splenic CD4^+^ or CD8^+^ T cells expressing PD-1 from 5 WT or 6 *Igha*^−/−^ mice determined by flow cytometry at steady state. (E) PD-L1-expression (MFI, mean fluorescence intensity) on splenic MZ B cells, FO B cells, total B-1 cells, B-1a cells, and B-1b cells from 5 WT or 6 *Igha*^−/−^ mice determined by flow cytometry at steady state. (F) ELISA of serum IgG1 to PPS 14, 21, or 28 days following i.p. immunization with Prevnar13 of 13 WT or *Igha*^−/−^ mice treated with anti-PD-1 mAb and of 11 WT or 12 *Igha-/-* mice treated with control isotype-matched mAb. Mice received 200 *μ*g i.p. of either anti-PD-1 or control mAb one day prior to immunization and one day after immunization, followed by injections every third day through day 19 post-immunization. (G) Frequency (%) of circulating anergic IgM^lo^ CD21^lo^ B cells within IgD^+^CD27^−^ naïve B cells from 17 healthy controls (HCs), 6 IGAD-R patients and 6 IGAD-NR patients. (H) ELISA of serum sCD14 from 18 HCs, 9 IGAD-R patients, and 6 IGAD-NR patients. Results summarize one (A, B, D, E, H), two (F), or 15 independent experiments (G), or show representative flow dot plots (C). Data are presented with mean GSVA score and 95% confidence interval (A) or mean and significance (D-H). Significance was determined using limma modeling (A), Mann- Whitney test (D, E, F) or Kruskal-Wallis test with Dunn’s correction for multiple comparisons (G, H). *p < 0.05, **p < 0.01. See also Figure S6.

Considering that the activation-induced receptor PD-1 triggers functional B cell (and T cell) unresponsiveness by transmitting immune inhibitory signals (Cambier et al., 2007; Okazaki and Honjo, 2007), we also evaluated whether IgA deficiency increased PD-1 and PD-ligand 1 (PD-L1) expression on systemic T and B cells, respectively. Compared to WT controls, *Igha*^−/−^ mice showed increased PD-1 expression by splenic CD4^+^ and CD8^+^ T cells as well as increased PD-L1 expression by splenic MZ and B-1 but not FO B cells (**Figure 6C-E**).

The involvement of PD-1 in the inhibition of vaccine-induced IgG1 in *Igha*^−/−^ mice was further evaluated with a blocking anti-PD-1 antibody. Consistent with earlier results, we found that *Igha*^−/−^ mice i.p. immunized with Prevnar13 and treated with a control isotype-matched antibody showed reduced serum IgG1 to PPS at day 21 and 28 compared to WT controls (**Figure 6F**). In contrast, immunized *Igha*^−/−^ mice treated with anti-PD-1 had as much serum IgG1 to PPS as WT mice treated with a control antibody (**Figure 6F**). Thus, anti-PD-1 treatment rescues the systemic impairment of the vaccine-specific IgG1 response caused by IgA deficiency.

Next, we determined whether, similar to B cells from *Igha*^−/−^ mice, B cells from IGAD patients exhibited phenotypic properties consistent with anergy, including lower IgM and CD21 expression (Duty et al., 2009; Isnardi et al., 2010). To address this question, the NYC cohort of adult IGAD patients was further stratified in IGAD responders (IGAD-Rs), who developed protective PPS-specific IgG responses following vaccination with Pneumovax23, and IGAD non-responders (IGAD-NRs), who did not (**Table S1**). Flow cytometry showed that adult IGAD-NRs had increased anergic IgM^low^CD21^low^ naïve B cells compared to healthy controls, while no difference was seen between IGAD-Rs and healthy controls (**Figure 6G**). This increase was associated with higher serum soluble CD14 (sCD14) (**Figure 6H**), a hallmark of systemic gut antigen translocation and immune activation in humans (Brenchley and Douek, 2012; Fadlallah et al., 2018).

Having shown that anergic B cells are increased in IGAD-NRs, we wondered whether these patients had additional B cell anomalies. Flow cytometry showed that the frequency of circulating total, naïve and MZ-like B cells was comparable in healthy controls, IGAD-Rs and IGAD-NRs (**Figure S6A** and **S6B**). However, the frequency of switched memory B cells, which typically emerge from the GC reaction (Cerutti et al., 2013), was decreased in IGAD-NRs compared to healthy controls (**Figure S6A** and **S6B**).

This finding was replicated in an additional Barcelona (BCN) cohort of adult IGAD patients, who also had a normal frequency of circulating total and MZ-like B cells and a decrease in switched memory B cells but an increased frequency of naive B cells (**Figure S6C** and **Table S2**). Furthermore, IgA^+^ PCs were decreased and IgG^+^ PCs showed a trend of increased frequency in IGAD compared to healthy controls. Of note, a second BCN cohort of IGAD children showed normal frequency of circulating total, naïve, MZ-like and switched memory B cells, but decreased frequency of IgA^+^ PCs and increased frequency of IgG^+^ PCs (**Figure S6D** and **Table S2**). Thus, in IGAD patients, B cell anomalies encompass IgA^+^ PC depletion in addition to IgG^+^ PC expansion, which echoes the increased commensal-specific IgG1 production observed in *Igha*^−/−^ mice. In summary, IgA may enhance systemic IgG responses to vaccines by limiting T and B cell expression of PD-1 and PD-L1, respectively. IgA could do so by restraining gut antigen translocation and subsequent B cell activation and anergy.

### IgA Increases Systemic Branched Chain Amino Acids Capable of Enhancing IgG Production

Given that gut metabolites regulate humoral immunity, exhibit antibody-limited penetration into host tissues and constrain PD-1 expression by T cells (Kim et al., 2016; Mager et al., 2020; Pedersen et al., 2016; Uchimura et al., 2018), we hypothesized that gut IgA interaction with commensals could further enhance systemic IgG responses to vaccines via metabolites. Accordingly, capillary electrophoresis time-of-flight mass spectrometry (CE-TOFMS) and liquid chromatography time-of-flight mass spectrometry (LC-TOFMS) showed less abundant branched-chain amino acids (BCAAs) such as valine, leucine and isoleucine in plasma from IGAD-NRs compared to healthy controls from the NYC cohort of adult IGAD patients (**Figure 7A** and **7B**). Plasma BCAAs were somewhat decreased also in *Igha*^−/−^ mice compared to WT controls, albeit not in a statistically significant manner (**Figure S7A** and **S7B**).

**Figure 7.**
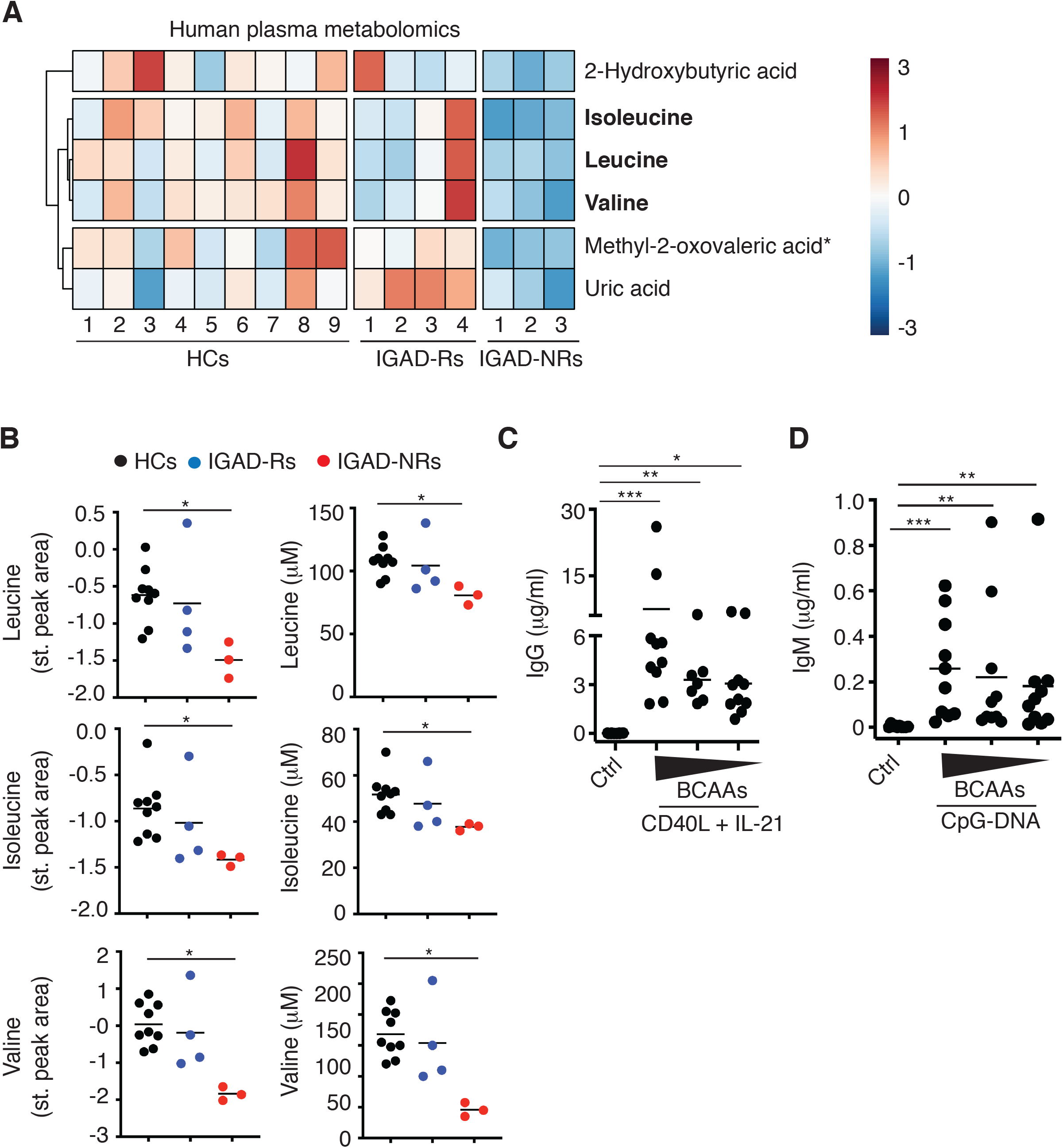
**IgA Increases Systemic BCAAs Capable of Enhancing IgG Production** (A) Heat map of significantly differentially abundant plasma metabolites from 9 HCs, 4 IGAD-R patients and 3 IGAD-NR patients. Data represent standardized peak areas and have been row mean-centered and row-normalized. Metabolites highlighted in bold are discussed in the text. SPA, standardized peak area. The positioning of the methyl group on methyl-2-oxvaleric acid was unable to be differentiated between attachment to either the third or fourth carbon of the molecule. (B) Plasma concentration of leucine, isoleucine and valine in donors described in A. Standardized peak area compared to internal standard and quantitative estimation (*μ*M) was performed by normalizing relative peak areas and comparing to standard curve. (C) ELISA of IgG secreted by B cells from 7-10 HCs upon exposure of peripheral blood mononuclear cells to medium alone (ctrl) or T cell-associated stimuli CD40L and IL-21 for 6 days in the presence of decreasing amounts of BCAAs. From left to right: complete BCAA-sufficient media, (120 mg/L BCAAs at a 2:5:5 ratio of L-valine, L-leucine, and L-isoleucine), BCAA-depleted media with one tenth as much BCAAs (12 mg/L BCAAs at the same ratio as above), and BCAA-deficient media with no BCAAs. (D) ELISA of IgM secreted by B cells from 10 HCs upon exposure of peripheral blood mononuclear cells for 6 days to medium alone (ctrl) or the TI ligand CpG-DNA in the presence of decreasing amounts of BCAAs as in (C). Metabolomics (A, B) were from one experiment. *In vitro* BCAA experiments (C, D) summarize two independent experiments involving peripheral blood mononuclear cells from 5 HCs each. Data are presented with mean and significance was determined through Kruskal-Wallis test with Dunn’s correction for multiple comparisons. *p < 0.05, **p < 0.01, ***p <0.001. See also Figure S7.

Although a few additional metabolites were differentially abundant in IGAD-NRs compared to healthy controls and IGAD-Rs, we opted to test the IgG-modulating potential of BCAAs due to their essential role in cell survival and to the key impact of the BCAA transporter CD98 on both TD and TI antibody responses (Cantor et al., 2009). Human B cell *in vitro* cultures followed by ELISA showed that, in the presence of decreasing amounts of BCAAs, TD signals such as those provided by CD40L and IL-21 or TI signals such as those provided by CpG-DNA induced progressively less IgG secretion (**Figure 7C** and **7D**).

The possible involvement of metabolites in the enhancement of systemic IgG responses by gut IgA was further explored by interrogating the impact of IgA deficiency on spermidine, an endogenous polyamine with known IgG enhancing properties in mice (Zhang et al., 2019). We found that spermidine was indeed decreased in the plasma from *Igha*^−/−^ mice compared to WT controls (**Figure S7B** and **S7C**). Thus, IgA could enhance systemic IgG responses to vaccines not only by limiting gut antigen translocation and PD-1 expression, but also by sustaining the peripheral abundance of metabolites with IgG-enhancing function, including BCAAs (**Figure S7D**).

## DISCUSSION

We have shown that gut IgA enhances peripheral IgG responses to TI or TD pneumococcal vaccines in both standard and gnotobiotic mouse models. By increasing the translocation of gut commensal antigens, IgA deficiency enhanced commensal-specific IgG production by B cells as well as expression of the negative feedback receptor PD-1 by T cells. Increased immune inhibitory signals from PD-1 impaired IgG production to pneumococcal vaccines by peripheral B cells, which exhibited a gene signature consistent with anergy. Moreover, in patients unresponsive to pneumococcal vaccines, IgA deficiency depleted peripheral gut metabolites such as BCAAs. These microbiota-regulated metabolites were found to amplify IgG production in human B cells exposed to TD or TI signals. Thus, gut IgA is functionally interconnected to peripheral IgG through multiple mechanisms involving the gut microbiota.

The gut microbiota enhances mucosal IgA and systemic IgG production to a broad spectrum of bacteria (Ansaldo et al., 2019; Fadlallah et al., 2019; Koch et al., 2016; Zeng et al., 2016). In this manner, commensals generate a pre-immune layer of humoral protection against pathogens aside from promoting gut homeostasis (Cullender et al., 2013; Rollenske et al., 2018). This dual function likely stems from the cross-reactivity of immunodominant antigens co-expressed by some commensals and pathogens (Cullender et al., 2013; Rollenske et al., 2018). As a result, a subset of patients with combined IgA and IgG depletion develop pathogen-driven systemic infections in addition to commensal-driven gut inflammation (Fadlallah et al., 2019; Uzzan et al., 2016).

The gut microbiota also sustains specific systemic IgG responses to vaccines (Hagan et al., 2019; Oh et al., 2014). This effect could involve IgA, because IgA is a mucosal antibody that extensively coats the gut microbiota (Magri et al., 2017; Palm et al., 2014). Accordingly, specific systemic IgG responses to pneumococcal vaccines are impaired in up to 30% of IGAD patients (Edwards et al., 2004; Lane and MacLennan, 1986). By taking advantage of multiple conventional and gnotobiotic mouse models, we found that IgA augmented IgG production to pneumococcal vaccines via gut commensal bacteria. Indeed, pIgR-deficient *Pigr*^−/−^ mice, which selectively lack mucosal secretory IgA, including gut SIgA (Johansen et al., 1999), exhibited an impairment of systemic IgG responses to pneumococcal vaccines similar to that of IgA-deficient *Igha*^−/−^ mice, which globally lack both systemic and mucosal IgAs (Harriman et al., 1999). In addition, we found that ex-GF mice reconstituted with gut bacteria from *Igha*^−/−^ mice or IGAD patients made less systemic IgG to vaccines compared to control ex-GF mice reconstituted with gut bacteria from IgA-sufficient mouse or human donors. Altogether, this evidence demonstrates that IgA is central to the induction of protective IgG responses by pneumococcal vaccines and exerts this effect through the gut microbiota. In this process, the gut microbiota and IgA likely form intertwined biological networks with interconnected IgG-enhancing functions.

Earlier seminal works explored the impact of the gut microbiota on IgG responses to vaccines such as the influenza vaccine (Hagan et al., 2019; Oh et al., 2014), but did not evaluate the contribution of IgA bound to intestinal microbes. Yet, this binding shapes both composition and function of gut commensal bacteria, including their metabolic properties (Fadlallah et al., 2018; Kawamoto et al., 2012; Macpherson et al., 2018; Nakajima et al., 2018). Furthermore, IgA binding to gut bacteria maximizes gut microbiota diversity, which is critical for optimal IgG responses to vaccines (Hagan et al., 2019; Kawamoto et al., 2012; Macpherson et al., 2018). Moreover, IgA constraints gut microbes to the intestinal lumen, which limits their interaction with the immune system (Chen et al., 2020; Macpherson et al., 2018).

Consistent with the key role of IgA in the control of intestinal bacteria, we found that IgA deficiency enriched the gut microbiota in *Lactobacillaceae* and caused abnormal translocation of *Lactobacillus (L.) murinus*, *L. reuteri* and *L. intestinalis* to the mesenteric adipose tissue. This last finding raises the possibility that intraluminal gut antigens undergo more translocation in mice lacking IgA. Such translocation could reflect impaired immune exclusion of Lactobacilli by gut IgA, which heavily coats Lactobacilli in the mouse intestine (Palm et al., 2014).

Our data indicate that the enhanced translocation of gut commensal antigens caused overstimulation of intestinal as well as systemic B, T and possibly myeloid cells in both *Igha*^−/−^ mice and IGAD patients. Indeed, in the absence of IgA, systemic B cells secreted abnormally larger amounts of IgG1 to translocated bacterial antigens in *Igha*^−/−^ mice. This increased IgG1 secretion likely involved a TD pathway that led to overstimulation of splenic FO B cells, a major B cell subset poised to undergo GC differentiation in response to antigenic stimulation. Consequently, GC B cells were remarkably increased in *Igha*^−/−^ mice along with GC-based TFH cells. In *Igha*^−/−^ mice, the lack of IgA also increased T cell expression of PD-1, a powerful immune inhibitory receptor that caused hyporesponsiveness of FO B cells to vaccine antigens. Accordingly, anti-PD-1 treatment rescued IgG1 responses to a pneumococcal vaccine in *Igha*^−/−^ mice. Similar to these mice, IGAD patients unresponsive to a pneumococcal vaccine showed increased B cells with anergic IgM^low^CD21^low^ phenotype. This finding was associated with that of augmented circulating sCD14, a soluble antigen released by myeloid cells as a result of hyperactivation by translocated gut bacteria antigens (Brenchley and Douek, 2012; Fadlallah et al., 2018). Thus, gut IgA may enhance vaccine-specific systemic IgG responses by restraining hyperactivation-induced and PD-1- mediated B cell hyporesponsiveness to pneumococcal vaccines.

In addition to constraining T cell expression of PD-1, intestinal IgA limited B cell expression of PD- L1, as this ligand increased in mice lacking IgA. Thus, gut IgA could attenuate IgG responses to pneumococcal vaccines by limiting PD-1 expression on T cells as well as PD-L1 expression on B cells, including MZ B cells. Unlike FO B cells, MZ B cells did not show transcriptional signatures reflecting abnormal activation in *Igha*^−/−^ mice. Yet, their increased PD-L1 expression may be central to PD-1 engagement on T cells and the ensuing initiation of immune inhibitory signals.

Interestingly, the gut microbiota has been recently shown to restrain PD-1 signaling via inosine, a metabolite that modulates T cells through the adenosine A2A receptor (Mager et al., 2020). This evidence implies that gut IgA might constrain PD-1 expression by supporting specific communities of inosine- releasing commensals. Should this be the case, gut IgA could enhance systemic IgG responses to pneumococcal vaccines by sustaining inosine production by intestinal commensal microbes in addition to constraining commensal antigen translocation.

Co-housing experiments showed that lateral transmission of a healthy gut microbiota into *Igha*^−/−^ mice could not rescue vaccine-specific IgG production. This finding suggests that IgA deficiency exerts a negative impact on systemic IgG responses through an early mechanism that may involve the vertical transmission of dysbiotic gut microbiota from the mother to the offspring. Consistent with this interpretation, *Igha*^−/−^ mice showed early hyperactivation of both mucosal and systemic immune responses, including steady-state overproduction of IgG1, together with impaired production of IgG3 to a CPS-mimicking TI immunogen. This evidence echoes recently published studies that demonstrate how gut IgA drives early post-natal changes of the gut microbiota and how gut microbes drive early post-natal immune development (Gomez de Aguero et al., 2016; Planer et al., 2016).

Our data imply that systemic IgG responses to vaccines may be irreversibly impaired soon after birth as a result of the early lack of gut IgA and the early perturbation of the gut microbiota. Such perturbation likely involves vertically transmitted microbial changes that exert dominant negative effects over healthy commensals newly acquired through lateral transmission and might entail the expansion of *Sutterella*, an IgA-degrading gut microbe (Kaakoush, 2020) that was more abundant in *Igha*^−/−^ mice compared to co-housed littermate controls. Hence, IgA deficiency may encompass phenotypic traits, such as impaired IgG responses to pneumococcal vaccines, through a mechanism involving the gut microbiota.

In agreement with this notion, gnotobiotic experiments indicated that ex-GF mice reconstituted with gut bacteria from IGAD patients made less intestinal IgA to commensals compared to control ex-GF mice reconstituted with gut bacteria from IgA-sufficient humans. Thus, gut IgA may amplify mucosal IgA responses to commensals in addition to systemic IgG responses to vaccines. This effect may not merely result from the intraluminal containment of commensal antigens and the restriction of IgA-degrading bacteria such as *Sutterella* (Kaakoush, 2020), but could further involve the intestinal retention of microbial consortia with IgA-inducing properties. Consistent with this possibility and similar to recently published data (Fadlallah et al., 2018), we found that IgA deficiency depleted *Ruminococcaceae* and *Lachnospiraceae* from the mouse intestinal microbiota. Of note, members of these large microbial families may amplify IgA responses to highly diverse and potentially beneficial mucus-embedded bacteria in the healthy gut from humans (Chen et al., 2020; Magri et al., 2017).

In addition to facilitating gut epithelial cell adherence of specific beneficial bacteria capable of excluding potentially harmful microbial competitors (Donaldson et al., 2018), gut IgA instigates competitive mucosal colonization by modulating the metabolic profile of certain commensals (Nakajima et al., 2018). Accordingly, we found that IgA deficiency caused systemic depletion of BCAAs in humans, a group of serum metabolites regulated by the gut microbiota (Pedersen et al., 2016). A trend towards BCAA depletion was also observed in mice, but did not reach statistical significance. Remarkably, BCAAs augment antibody responses to TD or TI immunogens by signaling through a multi-component transmembrane transporter called CD98, which is expressed by activated B cells (Cantor et al., 2009). Consistent with these published mouse data, *in vitro* cultures showed that BCAAs enhanced IgG production by human B cells exposed to TD or TI signals. Thus, IgA could sustain IgG responses to TD or TI pneumococcal vaccines through an additional mechanism involving BCAAs.

In the absence of IgA, mice showed depletion of the endogenous polyamine spermidine, which has been recently shown to increase systemic IgG responses to a TD immunogen such as chicken gammaglobulins (Zhang et al., 2019). This positive effect on post-immune IgG responses relies on the ability of spermidine to improve B cell function while constraining B cell senescence (Zhang et al., 2019). In summary, we show that gut IgA amplifies systemic IgG responses to pneumococcal vaccines through multiple mechanisms involving the gut microbiota. Aside from revealing an unexpected functional link between gut IgA and systemic IgG, our data indicate that gut IgA is a central component of interconnected intestinal IgG-inducing biological networks encompassing the microbiota. This implies that the potential contribution of gut IgA should be always considered in studies that evaluate the impact of intestinal microbes on local or systemic biological processes. Our findings also support the potential clinical benefit of oral IgA therapy in IGAD and common variable immunodeficiency patients with hyporesponsiveness to systemic vaccines, including pneumococcal vaccines.

## STAR**★**METHODS

### CONTACT FOR REAGENT AND RESOURCE SHARING

Further information and requests for resources and reagents should be directed to and will be fulfilled by Andrea Cerutti.

### EXPERIMENTAL MODEL AND SUBJECTS DETAILS

#### Human Blood and Stool Specimens

Human blood samples from the NYC cohort were collected from 42 age-and sex-matched HCs and IGAD patients at Icahn School of Medicine at Mount Sinai (**Table S1**). The age range was from 22 to 61 years (median 34 years) and 66.67% were females. Exclusion criteria for IGAD patients in the NYC cohort included serum IgA > 7 mg/dl, progression to common variable immunodeficiency, recent immunomodulatory treatments such as steroids or biologics targeting cytokines, antibiotics use within six weeks, or surgical procedures likely to affect humoral immunity such as splenectomy or intestinal resection. Responders (IGAD-Rs) and non-responders (IGAD-NRs) to pneumococcal vaccination were defined as IGAD individuals with a presence or absence, respectively, of protective serum IgG against more than 4 serotypes of 14 total after administration of Pneumovax23. Two IGAD patients determined to have an inadequate antibody response after receiving Pneumovax23 were immunized with Prevnar13 at a later date. However, neither of these two patients showed an increase in the number of protective IgG titers against pneumococcal serotypes in subsequent blood work. Compared to IGAD-Rs, a higher percentage of IGAD-NRs had a concomitant IgG2 or IgG4 subclasses deficiency (**Table S1**). Of note, IgG2 is the human equivalent of mouse IgG3 and mediates humoral immunity against PPS in humans (Briles et al., 1981; Siber et al., 1980).

A few years after the beginning of our study, the European Society for Immunodeficiencies revised the diagnostic criteria for selective IgA deficiency (https://esid.org/Working-Parties/Clinical-Working-Party/Resources/Diagnostic-criteria-for-PID2#Q7). According to these new criteria, a patient with selective IgA deficiency must have normal serum IgG and normal IgG responses to vaccination. While acknowledging these new diagnostic criteria, it may be worth considering that IgA deficient patients with impaired IgG responses to polysaccharides, including pneumococcal vaccines, remain poorly characterized. These patients, defined in our study as IGAD-NRs, may be part of a large spectrum of biological conditions in which IgA deficiency induces a progressive alteration of the IgG response. Selective IgA deficiency as defined by the new diagnostic criteria could stand at one extreme of this spectrum of IgA disorders, whereas common variable immunodeficiency progressing from selective IgA deficiency would stand at the opposite extreme (Aghamohammadi et al., 2008). IgA deficiency with impaired IgG responses to pneumococcal vaccines and/or combined IgG subclass deficiency would stand in between these two extremes of such heterogeneous spectrum of IgA disorders (Aghamohammadi et al., 2008; Edwards et al., 2004; Lane and MacLennan, 1986).

For the phenotypic study of B cells by flow cytometry, blood was collected in sodium-heparin blood collection tubes. For metabolomics analysis, blood from fasted patients was collected in BD Vacutainer K2EDTA blood collection tubes (Becton Dickinson). Human stool samples from the NYC cohort were collected from HCs and IGAD patients, shipped overnight on ice packs to the laboratory, and immediately stored at -80**°**C degrees. The Institutional Review Board of Icahn School of Medicine at Mount Sinai approved the use of blood and stool specimens. Human blood samples from the BCN cohort were collected from age- and sex-matched HCs and IGAD patients at Vall d’Hebrón Hospital (**Table S2**). The Ethical Committee of Clinical Research of Vall d’Hebrón Hospital approved the use of blood samples.

#### Mice

C57BL/6J (The Jackson Laboratory), *Igha*^−/−^ (Harriman *et al.,* 1999), and *Pigr*^−/−^ mice (Johansen *et al*., 1999) were bred in the animal facility of Icahn School of Medicine at Mount Sinai under specific pathogen free (SPF) conditions. *Igha*^−/−^ mice were obtained from Sergio Lira (Icahn School of Medicine at Mount Sinai), whereas *Pigr*^−/−^ mice were provided by Beth Garvy (University of Kentucky). WT and *Igha*^−/−^ breeders were set up from heterozygous *Igha*^+*/*−^ parents to control for microbiota and genetic background variability between strains. WT controls used in experiments came from these breeders except where pointed out. Similarly, WT and *Pigr*^−/−^ breeders were set up from heterozygous *Pigr*^+*/*−^ parents. Both sexes were used and mice were 4-16 weeks old for steady state experiments and 5-12 weeks old for immunization experiments. Since all mouse strains used in this manuscript mounted a specific IgM response upon immunizations, mice lacking induction of specific IgM were excluded from analysis due to presumed poor immunization. All animal experiments described in this study were approved by Institutional Animal Care and Use Committee (IACUC) of the Icahn School of Medicine at Mount Sinai and were performed in accordance with the approved guidelines for animal experimentation at the Icahn School of Medicine at Mount Sinai. GF C57BL/6J mice were bred in-house at the Mount Sinai Immunology Institute Gnotobiotic Facility in flexible vinyl isolators. To facilitate high-throughput studies in gnotobiotic mice, ‘‘out-of-the-isolator’’ gnotobiotic techniques were utilized (Faith et al., 2014). Shortly after weaning and under strict aseptic conditions, 28-42-days old GF mice were transferred to autoclaved filter-top cages outside of the breeding isolator and colonized with human or murine microbiota.

### METHOD DETAILS

#### Processing of Human and Mouse Specimens

##### Human Blood and Stool Samples

Fresh peripheral blood mononuclear cells (PBMCs) were obtained from sodium heparinized blood samples by separation on Ficoll Histopaque-1077 gradient (Sigma-Aldrich). Frozen human stool samples were pulverized under liquid nitrogen in a sterile hood. Approximately 500 mg from each donor was blended into a slurry (40-50 mg/mL) in pre-reduced bacterial LYBHIv4 culture medium (Sokol et al., 2008) containing 37 g/l Brain Heart Infusion (BHI) Broth (BD Biosciences), 5 g/l yeast extract (BD Biosciences), 1 g/l D-xylose, 1 g/l D-fructose, 1 g/l D-galactose, 1 g/l cellobiose, 1 g/l maltose, 1 g/l sucrose, 0.5 g/l N-acetylglucosamine, 0.5 g/l L-arabinose, 0.5 g/l L-cysteine, 1g/l malic acid, 2 g/l sodium sulfate, 0.05% Tween 80, 20 mg/mL menadione, 5 mg/l hemin, and 0.1 M MOPS (3-(N-morpholino)propanesulfonic acid) at pH 7.2. The slurries were passed through sterile 100-*μ*m strainers to remove large debris. To store for later gavage of gnotobiotic mice, slurries were diluted 1:20 in LYBHIv4 media with 15% glycerol and stored at -80**°**C.

##### Mouse Blood and Tissue Samples

Blood was collected from mice in ethylenediamine tetraacetic acid (EDTA) tubes pre-immunization (PI) and post-immunization at days 3, 7, 14, and 21 by submandibular vein puncture. Additional blood was collected in EDTA tubes from an orbital socket after sacrificing the animal. Plasma was isolated by incubating blood in EDTA tubes at room temperature for 20 minutes and spinning for 20 minutes at 15,000 G and collected into sterile microtubes for freezing at -80**°**C until further use. Murine tissue was collected after isoflurane-induced euthanasia followed by exsanguination and placed in Iscove’s Modified Dulbecco’s Medium (IMDM) supplemented with 2% heat-inactivated FBS (Life Techologies). Single cell suspensions were created from spleen and MLNs through tissue homogenization between frosted glass slides. PPs were incubated in dissociation buffer (10% FBS HBSS-/- with 5 mM EDTA and 15 mM HEPES) 20 minutes at 37**°**C before meshing with a 70-*μ*m cell-strainer (Fisher) and a syringe plunger. Splenic cell suspensions were treated with red blood cell lysis buffer (KD Medical) before staining for flow cytometry.

#### Colonization of GF Mice

Cecal contents from WT or *Igha*^−/−^ mice were dissolved in 5 ml sterile phosphate buffer solution (PBS) by vortexing. 200 *μ*L of this murine fecal slurry or 200 *μ*L of human fecal glycerol stock solution (see above) from HCs or IGAD patients was orally gavaged once into GF mice. Colonized mice were handled under aseptic conditions thereafter. In experiments involving human fecal bacteria, each human stool sample was transferred into several mice to control for transplantation variability. The involvement of multiple human donors allowed the control of biological variability. 12 or 18 GF recipient mice were reconstituted with fecal material from 4 HCs or 4 IGAD patients, respectively. All immunization experiments in reconstituted GF mice were performed at least 14 days after bacterial colonization.

#### Isolation and Identification of Translocated Bacteria

Spleen, MLNs, mesenteric adipose tissue and a small piece of small intestine (as a positive control) were aseptically dissected, weighed and collected into sterile PBS containing 0.5 g/ml reduced cysteine. Tissues were homogenized in 1 ml reduced PBS with sterile 3.5-mm beads using a bead beater (Bio-Spec) before clearing by centrifugation (600 × g, 3 minutes). Under strict anaerobic conditions, tissue homogenates were passed through 100 *μ*m filters and 100 *μ*l was inoculated onto reduced BBL^TM^ Chocolate II Agar (Becton Dickinson) and incubated under anaerobic conditions at 37°C for 72 hours. Colonies were counted and representative colonies were picked, expanded in liquid LYBHIv4 media (see above) and identified using MALDI-TOF mass spectrometry against a custom database following acetonitrile extraction (Bruker Biotyper) (Chudnovskiy et al., 2016).

#### Streptococcus pneumoniae culture

*Streptococcus pneumoniae* was streaked out on blood agar plates and incubated overnight at 37**°**C, after which 1 colony was picked with a sterile tip and inoculated in 5 ml pre-warmed BBL^TM^ Brain Heart Infusion Broth media (BD) and incubated at 37**°**C overnight. The overnight culture was then subcultured by transferring 500 *μ*l to 50 ml pre-warmed media (as above) and incubated 2-4 hours at 37**°**C until optical density (OD) 600 nm was approximately 0.3. *Streptococcus pneumoniae* was then made replication-deficient through incubation with 50 *μ*g/ml mitomycin-C (Sigma Aldrich) at 37**°**C for 1 h, and washed twice with PBS before immunization as described below.

#### Immunizations

Each mouse was i.v. or i.p. immunized with 2.87 *μ*g Pneumovax23 (Merck), 1.625 *μ*g Prevnar13 (Pfizer), 50 *μ*g TNP-aminoethilcarboxymethyl (AECM)-Ficoll (Biosearch Technologies), 50 *μ*g TNP-LPS (Biosearch Technologies). Each of these immunogens was diluted into 100-200 *μ*L sterile PBS before injection. 50 *μ*g NP-OVA-16 supplemented with 100 *μ*l Imject Alum Adjuvant according to manufacturer’s instructions (Thermo Scientific) was injected i.p. Replication-deficient *Streptococcus pneumoniae* prepared as above was i.v. injected at 2 × 10^7^ cells/mouse. For anti-PD-1 experiments, mice were treated with rat IgG2a mAb RMP1-14 to mouse PD-1 (Bio X Cell) or rat IgG2a mAb 2A3 as control (Bio X Cell). Mice received 200 *μ*g i.p. of either anti-PD-1 or control mAb in 200 *μ*l sterile dilution buffer (Bio X Cell) one day prior to Prevnar13 immunization and on days 1, 4, 7, 10, 13, 16, and 19 post-immunization.

#### Bacteria Isolation from Mouse Feces

Fecal pellets collected directly from mice or from the small intestine were homogenized in PBS (1 ml/0.1 g) by vortexing for 10 minutes at room temperature. Suspension was centrifuged twice at 2,000 rpm for 5 minutes at 4°C and supernatants collected. For bacterial flow cytometry assays, an aliquot was removed from this supernatant. After centrifugation at 8000 g for 10 minutes at 4°C, supernatant was collected to measure free fecal IgM and IgG1. Resulting bacterial pellets were used to determine microbiota-specific IgG1 (see ELISA).

#### Flow Cytometry and Cell Sorting

1-2 × 10^6^ cells per sample were resuspended in PBS with CD16/32 Fc Block (BD Biosciences) and Live/Dead staining (Invitrogen) for 10 minutes on ice. Cells were then washed with FACS buffer (PBS supplemented with 2% heat-inactivated FBS and 2 mM EDTA (Fisher)), and subsequently stained with appropriately diluted antibodies for 30 minutes on ice. For extracellular/intracellular IgG3 and IgG1, cells were blocked with 25% normal rat serum, stained for surface markers, including anti-IgG3-biotin followed by streptavidin-APC staining. After fixation and permeabilization (BD Cytofix/Cytoperm), cells were blocked again with 25% normal rat serum before addition of intracellular antibodies, including FITC-conjugated IgG3 diluted in BD Perm/Wash (BD Biosciences). For IgA-coated bacterial staining, bacteria isolated from fecal pellets were washed in staining buffer (1% BSA PBS w/v) and blocked for 20 minutes on ice with staining buffer supplemented with 20% normal rat serum (Invitrogen) prior to staining with anti-IgA antibody 1:100 (final 1:200) for 30 minutes (Palm et al., 2014). To quantify serum IgG1 specific to commensals, bacteria were optionally incubated with matched mouse serum diluted 1:50 in staining buffer and incubated on ice for 30 minutes. After washing with staining buffer, samples were stained with a biotinylated anti-IgG1 antibody (1:25) for 30 minutes on ice before another wash and staining with streptavidin-APC. After washing, bacteria were resuspended in a saline buffer including 0.9% NaCl and 10 mM HEPES and supplemented with 1.25 nM SYTO Green Fluorescent Nucleic Acid Stain (Life Technologies). Cells were acquired with LSR Fortessa or LSR II (BD Biosciences) and data were further analyzed by FlowJo V10 software (TreeStar).

For cell sorting, B cells were purified using a Miltenyi Pan B cell isolation Kit II (Cat. 130-104-443), followed by FACS staining for sorting. Cells were resuspended in PBS with CD16/32 Fc Block and Live/Dead staining (Invitrogen) before staining with antibodies to surface markers. Live CD11c^−^Ly6G^−^CD3^−^ B220^+^AA4.1^−^ CD21^high^CD23^lo^ MZ B cells and CD21^+^CD23^+^ FO B cells were FACS-sorted (Influx BD) with Influx (BD Biosciences). The purity of sorted cells was consistently >95%.

#### ELISA

For total Ig subclass quantification, Immulon 4 HBX 96-well plates (ThermoFisher Scientific) were coated with a primary antibody diluted in UltraCoat ELISA Coating Buffer (Leinco Technologies) at 50 *μ*l per well overnight at 4**°**C. Antigen-specific Ig subclass quantification was determined by coating Immulon 4HBX 96-well plates overnight at 4**°**C with 50 *μ*l/well of the following antigens diluted in PBS at 5 *μ*g/mL: TNP-bovine serum antigen (BSA), CPS9 or CPS14 from *Streptococcus pneumoniae*, *Staphylococcus aureus*-derived lipoteichoic acid, *Salmonella typhimurium*-derived LPS, or mitomycin-C-inactivated whole *Streptococcus pneumonia* (4 ×10^6^ bacteria/mL). For detection of high- and low-affinity

NP-specific antibodies, plates were coated with NP-BSA ratio 7 (NP7) or NP-BSA ratio 23 (NP23), respectively, (Biosearch Technologies) at 10 *μ*g/mL. Coated plates were then washed with 0.1% polysorbate 20 (Fisher) in PBS and blocked with milk powder or 1% BSA in PBS for 2 hours at room temperature. After washing, serum or fecal supernatant serially diluted in ELISA dilution buffer (1% BSA 0.1% polysorbate 20 PBS) was added to each well and incubated overnight at 4**°**C. After additional washing, plates were incubated for 2 hours at room temperature with a horseradish peroxidase (HRP)- conjugated secondary antibody (Southern Biotech) diluted in ELISA dilution buffer. After further washing, plates were developed with two-component tetramethylbenzidine HRP substrate (Seracare Life Sciences) and stopped with an equal volume of 1M H2SO4. OD values were determined at 450-nm wavelength on Epoch microplate spectrophotometer (BioTek). Mouse IL-4 was measured using an IL-4 mouse uncoated ELISA Kit (Invitrogen). Human sCD14 was measured through Human CD14 DuoSet ELISA kit following manufacturer’s instructions (R&D Systems). For microbiota-specific IgG1 quantification, bacteria pellets from colonic fecal content was resuspended in 500 *μ*l PBS and the resulting suspension was frozen in dry-ice and then thawed in a 37°C water bath for five cycles (Hepworth et al., 2013). The resulting lysates were centrifuged at 8000 g for 10 min and supernatants were collected as commensal antigen lysates. Lysate protein concentration was measured with Nanodrop. Immulon 4 HBX 96-well plates (Thermo Scientific) were coated with 5 *μ*g/mL lysate protein in PBS and incubated overnight at 4°C. In each mouse, commensal-specific titers were determined as the difference in OD values between samples with or without paired serum. When not expressed in units of optical density (OD), the relative concentration of antigen-specific antibodies in serum from immunized mice was expressed in arbitrary units (AU) calculated from a standard curve obtained with serum from previously-immunized mice with high titers of specific antibodies run contemporaneously. An appropriate control serum was used as standard to quantify total antibody concentrations in mice (Bethyl Laboratories, Inc.).

#### RNA-Seq Analysis

RNA was extracted from sorted splenic murine MZ and FO B cells using QIAshredder and RNeasy Plus Micro Kit (Qiagen). Library preparation and sequencing were performed by a specialized company (GENEWIZ).

##### Standard Quality Control and Mapping of Sequence Reads

The fastq files generated from Illumina RNA-Seq were preprocessed and quality controlled using the CLC Genomics Workbench Version 9.0.1. The raw reads were trimmed and filtered to remove possible adapter sequences and low-quality nucleotides at the ends. After trimming, sequence reads shorter than 50 nucleotides were discarded according to the following parameters: phred score > Q30, base error probability score > 0.05, and base ambiguities < 2, read-length < 50 bp. Trimmed reads were aligned to the *Mus musculus* genome, and count data was generated using CLC Genomics Workbench Version 9.0.1.

##### Differential Expression Analysis

Statistical analysis was carried out using R-language (R-project.org) and packages available through the Bioconductor project (www.bioconductor.org). Normalization factors to scale RNA-seq library size were calculated using TMM method (Robinson and Oshlack, 2010; Wanigasuriya et al., 2020) and converted to counts per million (CPM) reads using the cpm function (edgeR package). Gene transcripts with a CPM >1 in at least one sample were kept for further analysis. The remaining data was then transformed using *voom* (Law et al., 2014). Gene/transcript-expression profiles of non-Igh genes were compared between B cells from *Igha*^−/−^ and WT mice using linear mixed-effect models with cell-type (MZ vs FO B cells) and mouse strain (WT vs *Igha*^−/−^ mice) as fixed factors. A random intercept was used for each parent mouse the B cells were derived from. Models were fitted using the *limma* package framework and hypothesis of interest were tested via contrasts using the moderated t-test (Smith, 2004). P-values were corrected for multiple hypothesis testing and are presented with false discovery rate (FDR) values. Gene and transcripts annotation was carried out with Biomart (Durinck et al., 2005; Durinck et al., 2009) using the *mmusculus_gene_ensembl* dataset from 01/07/2019.

##### Gene Set and Gene Pathway Analysis

Over-representation analysis (ORA) and gene-set enrichment analysis (GSEA) were performed with the HALLMARK and C2CP collections from the MolSigDB database (http://software.broadinstitute.org/gsea/msigdb/index.jsp) as well as custom gene sets. Custom gene sets representing GC B cell differentiation and PC differentiation were defined using publicly available GSE60927 data (Shi et al., 2015) as genes up-regulated (fold change (FCH) >2, FDR <0.05) when comparing GC B cells vs FO B cells (GC B cell differentiation), and the top-50 down-regulated (FCH >1.5, FDR <0.05) genes when comparing bone marrow PCs vs splenic plasmablasts (this differentially expressed gene set was defined as PC differentiation). A gene-set representing B cell anergy was defined as up-regulated genes (FCH >2, FDR <0.05) when comparing anergic MD4ML5 cells vs naïve non-anergic MD4 cells (control) using data previously published (Sabouri et al., 2016). Gene-set variation analysis (GSVA) (Hänzelmann et al., 2013) was used to calculate gene-sets activity per sample for proliferation (Hallmark_G2M_checkpoint) and custom gene-sets. Statistical analysis of GSVA scores was performed using the same methods as gene-expression (limma package framework). Sequence data files (fastq) are stored in the public Sequence Read Archive (SRA) under project number GSE173361.

#### 16S rRNA Gene Sequencing and Analysis

DNA extraction, library preparation, and 16S rDNA gene amplicon sequencing of colonic fecal bacteria from co-housed littermate WT and *Igha*^−/−^ mice were outsourced (GENEWIZ). Barcoded amplicons were subjected to multiplexed sequencing (paired-end 250 nucleotide reads) on a MiSeq instrument (Illumina) with the 500 cycle v2 kit (2 x 250 bp). Paired-end reads were filtered (Phred > 19) and merged using the fastq-join algorithm. Data analysis was performed using QIIME with default parameters unless otherwise noted, and performed as previously described (Shapiro et al., 2021). Briefly, raw sequencing data were demultiplexed using unique barcodes assigned to each sample, and low quality reads discarded from downstream analyses. Remaining reads were then clustered into Operational Taxonomic Units (OTUs) with a 97% similarity threshold, and using the gg_13_5 release from Greengenes v13-8 as a reference set to assign taxonomy to each OTU (DeSantis et al., 2006; McDonald et al., 2012). The default open reference Qiime pipeline for Ilumina reads was also used (Caporaso et al., 2010). Taxonomy was assigned using default uclust algorithm parameters (Edgar, 2010). Chimeras were identified and removed using ChimeraSlayer software (Haas et al., 2011). There were no significant differences in sequencing depth per group. Processed spleen and MLN samples from GF mice were used to identify putative contaminant OTUs, which were then removed from all the samples. Next, an ‘abundance-filtered data set’ was generated by selecting OTUs with >0.1% relative abundance in each sample. This OTU table was then rarefied to the minimum sample’s depth (1570 reads). Scaled abundance heatmap was generated after selecting the 20 most important bacterial species highlighting the competing phenotypes as defined by the RandomForest algorithm (Liaw & Wiener 2002) from Qiime (ntree=50000,e=loo). Sequence data files (fastq) are stored in the public Sequence Read Archive (SRA) under project number PRJNA722766.

#### Metabolomics

Sera were collected from HCs, IGAD patients, SPF WT mice or SPF *Igha*^−/−^ mice as described above. HCs and IGAD patients were asked to fast overnight, whereas food was withheld from mice overnight prior to collection. Collected serum was frozen at −80**°**C and shipped to Human Metabolome Technologies for further analysis. Metabolome analysis was performed using both capillary electrophoresis time-of-flight mass spectrometry (CE-TOFMS) and liquid chromatography time-of-flight mass spectrometry (LC-TOFMS) in two modes for cationic and anionic metabolites. For CE-TOFMS, each 50 μL sample was mixed with 450 μL of 10 μM methanol-containing internal standards and mixed. Then, 500 μl chloroform and 200 μl Milli-Q water were added, mixed thoroughly and centrifuged at 2,300 × g, 4°C for 5 minutes. The water layer (400 μL × 1) was filtrated through a 5-kDa cut-off ULTRAFREE-MC-PLHCC filter (Human Metabolome Technologies) to remove macromolecules. The filtrate was centrifugally concentrated and resuspended in 50 μL of ultrapure water immediately before the measurement. For LC-TOFMS, each 500 μl (human) or 300 μl (mouse) sample was mixed with either 1,500 μl (human) or 900 μl (mouse) of 1% formic acid in acetonitrile (v/v) containing 6 μM internal standards and centrifuged at 2,300 x g, 4°C for 5 minutes. Then, the supernatant was filtrated by using a Hybrid SPE phospholipid 55261-U column (Supelco) to remove phospholipids. The filtrate was desiccated and resuspended in either 200 μl (human) or 120 μl (mouse) of 50% isopropanol in Milli-Q water (v/v) immediately before the measurement.

Peaks detected in CE-TOFMS or LC-TOFMS analysis were extracted using an automatic MasterHands ver. 2.17.1.11 integration software (Keio University) in order to obtain peak information, including m/z, migration time (MT) in CE, retention time (RT) in LC, and peak area. The peak area was then converted to relative peak area by dividing the Metabolite Peak Area by the Internal Standard Peak Area. The peak detection limit was determined based on signal-noise ratio; S/N = 3. Putative metabolites were then assigned from HMT’s standard library and Known-Unknown peak †4 library on the basis of m/z and MT or RT. The tolerance was ± 0.5 minutes in MT/RT and ±10 ppm. If several peaks were assigned the same candidate, the candidate was given the branch number. Absolute quantification was performed in 110 metabolites, including glycolytic and tricarboxylic-acid-cycle-intermediates, amino acids, and nucleic acids. All the metabolite concentrations were calculated by normalizing the peak area of each metabolite with respect to the area of the internal standard and by using standard curves, which were obtained by single-point (100 μM) calibrations. Standardized relative peak area data were used to identify metabolites showing statistically significant differences between defined group phenotypes (Kruskal Wallis with Dunn’s correction for human data and Mann-Whitney for mouse data, p value <0.05). Data represent standardized values of relative area in detected peaks where values below the limit of detection were assigned a value of epsilon(2^-52). Heat maps show standardized relative peak areas of significantly differentially abundant metabolites and have been row mean-centered and row-normalized. Heat maps were generated using the *gplots* R-package.

#### Mouse B Cell Cultures

A single cell suspension of splenocytes was prepared as above and purified for B cells using either MACs Mouse B cell Isolation Kit (Miltenyi Biotech) or EasySep Mouse B cell Isolation Kit (Stemcell Technologies). Cells were plated at 0.2 × 10^6^ cells/ml in 96-well round bottom plates in 200 μl RPMI medium containing 10% FBS, 1% penicillin-streptomycin (Life Sciences) and 55 μM 2-mercaptoethanol and stimulated with 1 μg/ml anti-CD40 (Biolegend) and 100 ng/ml IL-4 (R&D Systems), or 10 μg/ml LPS (Invivogen) for 6 days. Supernatant was collected on day 6 for ELISA and cells were harvested for flow cytometry.

#### Human B Cell Cultures

3 × 10^5^ PBMCs (200 *μ*l/well) were cultured in 96-well round bottom plates and stimulated with 500 ng/ml trimeric CD40L (Enzo Life Sciences) and 100 ng/ml IL-21 (Peprotech), or 500 ng/ml CpG-DNA (Invivogen) for 7 days. RPMI culture medium containing 10% FBS and 1% penicillin-streptomycin (Life Sciences) was prepared without BCAAs, with BCAAs at optimal concentrations in complete BCAA- sufficient media, (120 mg/L BCAAs at a 2:5:5 ratio of L-valine (0.17 nmol/L), L-leucine (0.38 nmol/L), and L-isoleucine (0.38 nmol/L), BCAA-depleted media with one tenth as much BCAAs (12 mg/L BCAAs at the same ratio as above). Supernatant was collected at day 7 and analyzed for immunoglobulin titers via ELISA.

#### Histology and Immunofluorescence

Colon and small intestine from WT or *Igha*^−/−^ mice were incubated for 4h in Methanol-Carnoy’s fixative at room temperature, then immersed in 100% ethanol at 4°C for 1h prior incubation with xylene solution at room temperature. Tissue was then embedded in paraffin solution and 4-*μ*m sections were cut before staining with hematoxylin and eosin at Mount Sinai’s Biorepository and Pathology CoRE service. Slides were analyzed with an Aperio AT2 (Leica) and scored by a pathologist from Mount Sinai Hospital. For immunofluorescence assays, whole spleens were excised from WT or *Igha*^−/−^ mice 30 minutes or 3 h following i.v. immunization with TNP-Ficoll, embedded in Optimal Cutting Temperature Compound (TissueTek), and frozen at −80**°**C. Sections were prepared with Leica CM1950 Cryostat and fixed with 4% paraformaldehyde for 15 minutes on ice after washing with PBS and permeabilized with Triton X-100 for an additional 15 minutes before blocking. Tissue was then incubated with unconjugated TNP-specific rat antibody (clone 2A3), followed by a488-conjugated anti-rat antibody, biotinylated anti-MOMA-1, APC-conjugated streptavidin and PE-eFluor610-conjugated anti-B220 antibody. Slides imaged using a DM600 Leica microscope and analyzed at Mount Sinai Microscopy Core and Advanced Bioimaging Center.

#### Quantification and Statistical Analysis

Statistical analysis was performed using Prism version 9.0 (GraphPad). Comparisons between two groups were determined using either Student’s t test when data followed a normal distribution or Mann-Whitney U test when data were not normally distributed. Normal/Gaussian distribution was determined by a D’Agostino and Pearson normality test. For multiple comparisons, a Kruskal-Wallis test with Dunn’s correction was used. Significant outliers were determined using Grubb’s test with an alpha of 0.05 and excluded from analysis. A p value < 0.05 was considered significant. P values are indicated on plots and in figure legends. (* p < 0.05, ** p < 0.01, *** p < 0.001).

## SUPPLEMENTAL INFORMATION

Supplemental Information includes seven figures and two tables.

## AUTHOR CONTRIBUTIONS

C.G. designed and performed research with both mouse models and human samples, discussed data, prepared figures, and helped editing the manuscript; E.K.G. performed research with mouse models, analyzed and discussed data, prepared figures, and helped to write and edit the manuscript; D.B.M. performed research with mouse models and human samples, analyzed and discussed data, and prepared figures; P.J.M. performed research with human samples, and analyzed and discussed data; G.M. performed research with human samples, analyzed and discussed data, and helped to edit the manuscript; G.J.B. helped with experiments involving GF and gnotobiotic mice; S.T.V. and R.T-P. performed research with human samples; L.T. and M.P. provided support with bioinformatics; P.K.V. helped with mouse breeding and performing assays; A.M.N., M.G.P., M.M.G. and R.D.-C. discussed data and organized the logistics of the BCN cohort; J.C.C. and S.M. discussed data and helped to edit the manuscript; M.S-F. provided support in bioinformatics and discussed data; J.J.F. profiled the mouse intestinal microbiota, helped with experiments involving GF and gnotobiotic mice, and edited the manuscript; C.C.-R. organized the logistics of the NYC cohort, designed research, analyzed and discussed data, and helped to edit the manuscript; A.C. designed research, analyzed and discussed data, and wrote the manuscript.

## ACKNOWLEDGMENTS

Supported by US National Institutes of Health grants P01 AI61093 (to A.C. and C.C-R.), R01 DK123749 (to A.C., J.J.F. and S.M.), R01 DK114038 (to J.C.) and K23 AI137183 (to P.J.M.); by Ministerio de Ciencia, Innovación y Universidades grant RTI2018-093894-B-I00 and European Advanced Grant ERC- 2011-ADG-20110310 (to A.C.); and by the Institute of Health Carlos III-Miguel Servet research program (to G.M.).

