## Supplemental Tables and Figures for "Gut IgA Enhances Systemic IgG Responses to Pneumococcal Vaccines Through the Commensal Microbiota"

### Supplemental Tables S1 and S2

### Supplemental Figures and Legends S1 – S7

**Table S1 (Related to Figures 3, 6, 7 and S6). NYC Cohort of IGAD Patients**

| **Relevant phenotypic traits** | **HCs** (n = 27) | **IGAD-Rs** (n = 10) | **IGAD-NRs** (n = 6) |
|---|---|---|---|
| **Gender** (% female) | 66.67% | 66.67% | 66.67% |
| **Age,** years (median [IQR]) | 33 [10] | 33 [13] | 40 [3] |
| **IgG-targeted Pneumococcal serovars post-Pneumovax23** (median of 14) | N/A | 11 [4] | 2 [3] |
| **Serum antibody titers, mg/dL** (median [IQR])[a] | | | |
| IgA | 196 [105] | <7 | <7 |
| IgM | 110 [51] | 84 [51] | 72 [116] |
| IgG | 1170 [268] | 1111 [455] | 812 [322] |
| IgG1 | 644 [117] | 729 [99] | 621 [264] |
| IgG2[b] | 381 [119] | 312 [173] | 125 [100] |
| IgG3 | 39 [39] | 61 [40] | 34 [120] |
| IgG4[c] | 29 [36] | 12 [61] | 1 [0] |
| **Recurrent infections,** *n* (%) | | | |
| Bronchitis | N/A | 5 (62.5) | 4 (80) |
| Otitis | N/A | 1 (12.5) | 1 (20) |
| Pneumonia | N/A | 5 (62.5) | 2 (40) |
| Sinusitis | N/A | 7 (87.5) | 3 (60) |

IQR, interquartile range

[a]Ig classes: HCs, n = 11; IGAD-Rs, n = 8; IGAD-NRs, n = 6.
IgG subclasses: HCs, n = 8; IGAD-Rs, n = 8; IGAD-NRs, n = 5.
[b]Of patients with IgG subclass data, 1 IGAD-R and 3 IGAD-NR had IgG2 titers below the sufficiency cut-off of 124 mg/dL.
[c]Of patients with IgG subclass data, 3 IGAD-R and 4 IGAD-NR had IgG4 titers below the sufficiency cut-off of 124 mg/dL

Inclusions per assay:
Flow cytometry: HCs, n = 17; IGAD-Rs, n = 6; IGAD-NRs, n = 6.
sCD14 quantification: HCs, n = 18; IGAD-Rs, n = 9; IGAD-NRs, n = 6.
Metabolomics: HCs, n = 9; IGAD-Rs, n = 4; IGAD-NRs, n = 3.
GF mouse reconstitution: HCs, n = 4; IGAD-Rs, n = 2; IGAD-NRs, n = 2.
*In vitro* assays: HCs, n = 10.

**Table S2 (Related to Figure S6). BCN Cohort of IGAD patients**

| **Relevant phenotypic traits** | **Pediatric HCs** (n = 11) | **Pediatric IGAD patients** (n = 14) | **Adult IGAD patients** (n = 17) |
|---|---|---|---|
| **Gender** (% female) | 45.45% | 42.85% | 64.71% |
| **Age**, years (median [IQR]) | 8 [7] | 10 [5] | 56 [19] |
| **Serum antibody titers,** mg/dL (median [IQR])[a] | | | |
| IgA | 123 [47] | <10 | <10 |
| IgM | 117 [66] | 79 [34] | 95 [82] |
| IgG | 1138 [143] | 1235 [243] | 1278 [555] |
| IgG1 | 813 [26] | 999 [210] | 807 [297] |
| IgG2 | 312 [58] | 335 [271] | 300 [256] |
| IgG3 | 74 [2] | 43 [14] | 44 [41] |
| IgG4 | 17 [4] | 35 [98] | N/A |

IQR, Interquartile range.

[a]Ig classes: pediatric healthy controls, n = 7; pediatric IGAD patients, n = 14; adult IGAD patients, n = 17.

IgG subclasses: pediatric healthy controls, n = 2; pediatric IGAD patients, n = 12; adult IGAD patients, n = 17.

Adult HCs, n = 23; relevant phenotypic traits were not available.

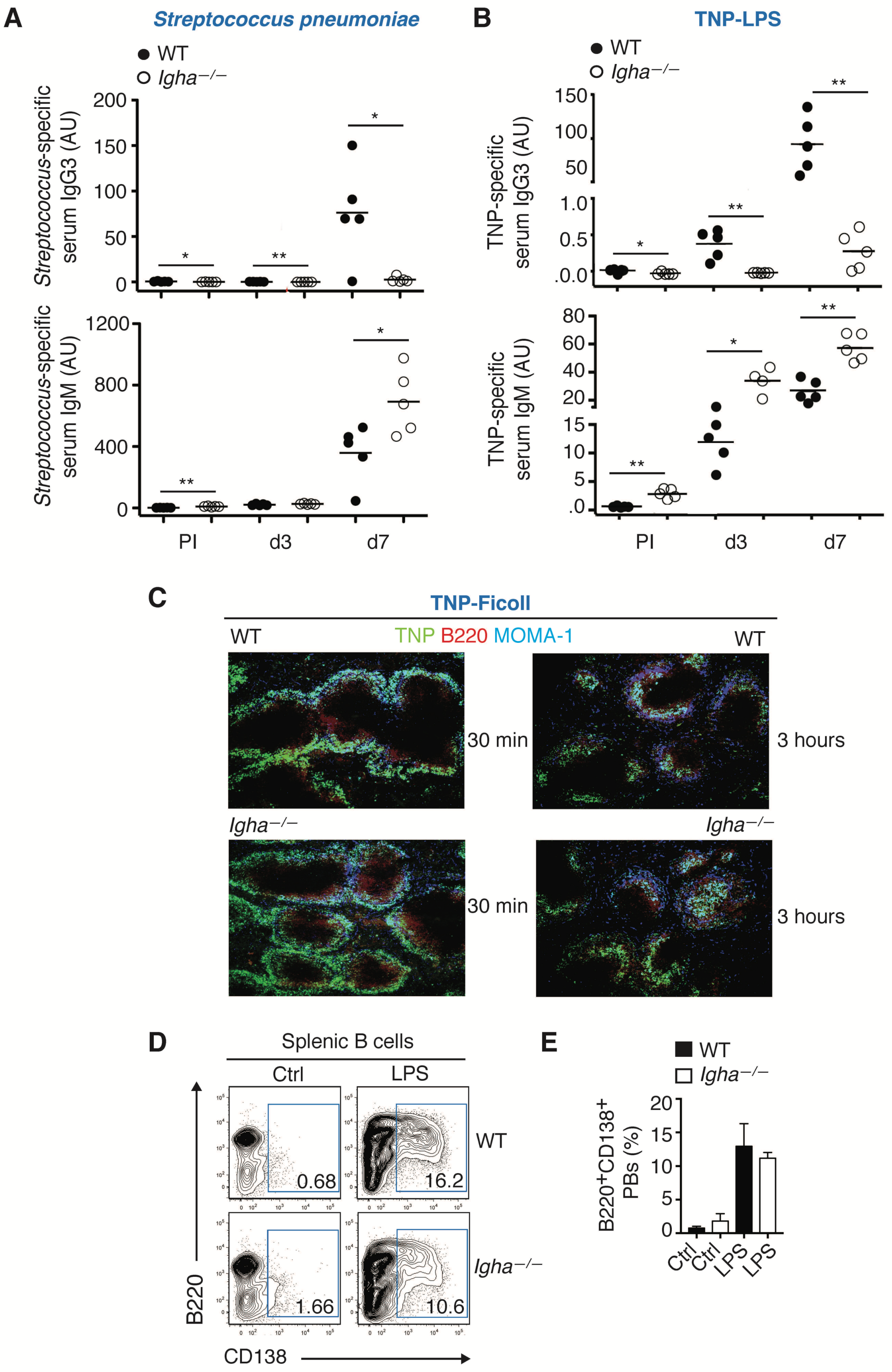


**Figure S1 (Related to Figure 1). IgA Enhances Systemic IgG Responses to TI Immunizations**

(A) ELISA of serum IgM and IgG3 to whole mitomycin C-inactivated *Streptococcus pneumoniae* from 5 WT or *Igha*$^{-/-}$ mice prior to immunization (PI) or 3 and 7 days following i.v. immunization with mitomycin C-treated non-replicating *Streptococcus pneumoniae*. WT mice in this experiment were purchased from Jackson Laboratories.

(B) ELISA of serum IgM and IgG3 to TNP from 5 WT and 4-5 *Igha*$^{-/-}$ mice PI or 3 and 7 days after i.v. immunization with TNP-LPS.

(C) IFA of the TNP hapten (green), the surface B cell molecule B220 (red), and the surface metallophilic macrophage molecule MOMA-1 (blue) from spleen sections of representative WT or *Igha*$^{-/-}$ mice 30 minutes (left) or 3 hours (right) following i.v. immunization with TNP-Ficoll.

(D) Flow cytometry of B220 and CD138 molecules on purified splenic B cells from representative WT or *Igha*$^{-/-}$ mice following exposure to medium alone (Ctrl) or LPS for 6 days. B220$^{+}$CD138$^{+}$ cells correspond to PBs. Numbers indicate PB frequency (% of live cells).

(E) Summary of the frequency (% of live) of B220$^{+}$CD138$^{+}$ PBs obtained from purified splenic B cells isolated from 6 WT or *Igha*$^{-/-}$ mice and cultured as in (D).

Results summarize one (A, B) or three (E) experiments involving 2 WT or *Igha*$^{-/-}$ mice or show representative images from an experiment involving 4 WT or *Igha*$^{-/-}$ mice (C) or representative plots from mice summarized in Figure S1E (D). Data are presented with mean (A, B) or mean ± s.e.m. (E); significance was determined using a two-tailed unpaired Mann-Whitney test; *P < 0.05, **P < 0.01

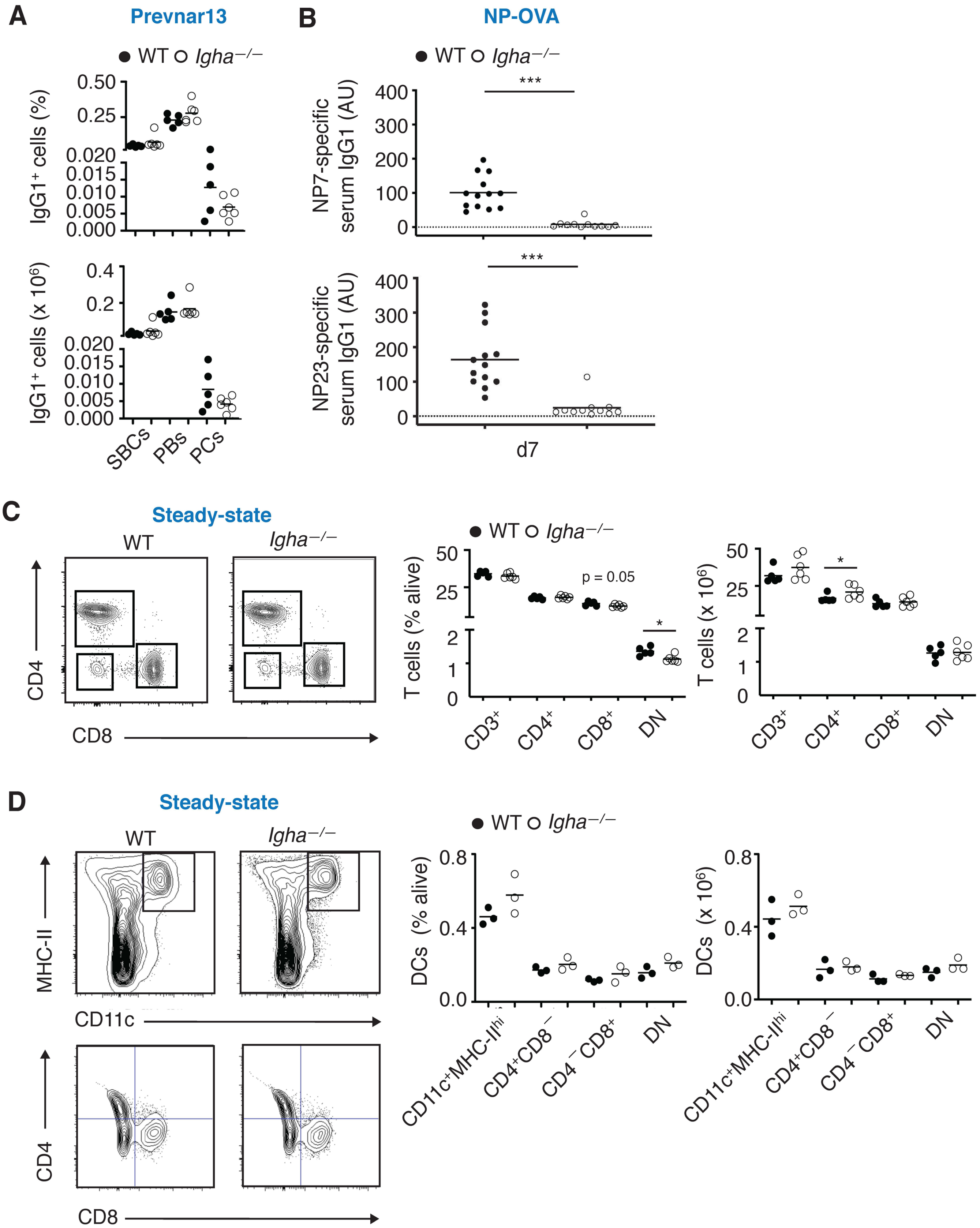


**Figure S2 (Related to Figure 2). IgA Increases Systemic IgG responses to TD Immunizations**

(A) Flow cytometry of intracellular (ic) IgG1 and extracellular (ec) IgG1 expression by splenic switched $ecIgG1^{+}icIgG1^{-}$ B cells (SBCs), $ecIgG1^{+}icIgG1^{+}$ PBs and $ecIgG^{lo}icIgG^{+}$ PCs from 5 WT or 7 *Igha*$^{-/-}$ mice 28 days following i.v. immunization with Prevnar13. Frequency (% of $B220^{+}$ cells, top) and absolute number (bottom) of SBCs, PBs and PCs are indicated.

(B) ELISA of serum high-affinity IgG1 to NP7-BSA and low-affinity IgG1 to NP23-BSA in 13 WT or 11 *Igha*$^{-/-}$ mice 7 days following i.p. immunization with NP15-OVA and alum. Data from individual mice are identical to d7 data from Figure 2C.

(C) Flow cytometry of total $CD3^{+}$ T cells, $CD4^{+}$, $CD8^{+}$ and $CD4^{-}CD8^{-}$ (double negative, DN) T cell subsets from the spleen of 5 WT or 6 *Igha*$^{-/-}$ mice at steady state. Representative flow cytometry contour plot (left) pre-gated on live $CD3^{+}$ T cells, frequency of T cell subsets (% live, middle), and absolute cell numbers (right) are indicated.

(D) Flow cytometry of total $CD11c^{+}MHC\text{-}II^{high}$ dendritic cells (DCs) as well as $CD4^{+}CD8^{-}$, $CD4^{-}CD8^{+}$ and $CD4^{-}CD8^{-}$ (DN) DC subsets from the spleen of 3 WT or *Igha*$^{-/-}$ mice at steady state. Representative flow cytometry contour plots (left), frequency (% live) and absolute numbers are indicated. DCs were first gated on $CD3^{-}B220^{-}$ cells. WT mice in this experiment were purchased from Jackson Laboratories.

Results summarize 1 (C and D right) or 2 (A, B) experiments or show representative flow cytometry plots (C and D left). Data are presented with mean; significance was determined using a two-tailed unpaired Mann-Whitney test. *$p < 0.05$, ***$p < 0.001$.

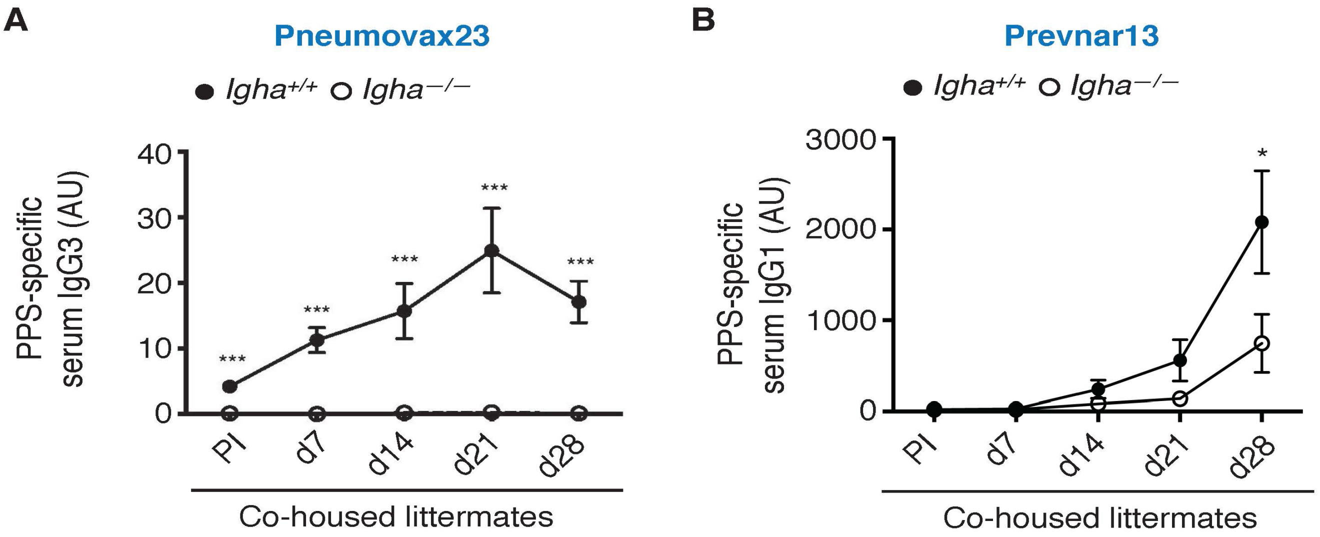


**Figure S3 (Related to Figure 3). IgA Helps Systemic IgG Responses to Vaccines through Gut Commensals**

(A, B) ELISA of serum IgG1 to PPS from *Igha*$^{+/+}$ or *Igha*$^{-/-}$ co-housed littermate mice from *Igha*+/- parents prior to immunization (PI) and 7, 14, 21, or 28 days following i.p. immunization with either Pneumovax23 (A) or Prevnar13 (B). Results from Pneumovax23 immunizations summarize one experiment with 9 *Igha*$^{+/+}$ WT mice and 5 *Igha*$^{-/-}$ mice (A), whereas results from Prevnar13 immunizations summarize 2 experiments with 7 *Igha*$^{+/+}$ and 7-8 *Igha*$^{-/-}$ mice (B).

Data are presented as mean ± s.e.m.; significance was determined using two-tailed unpaired Mann-Whitney test; *p <0.05, ***p <0.001.

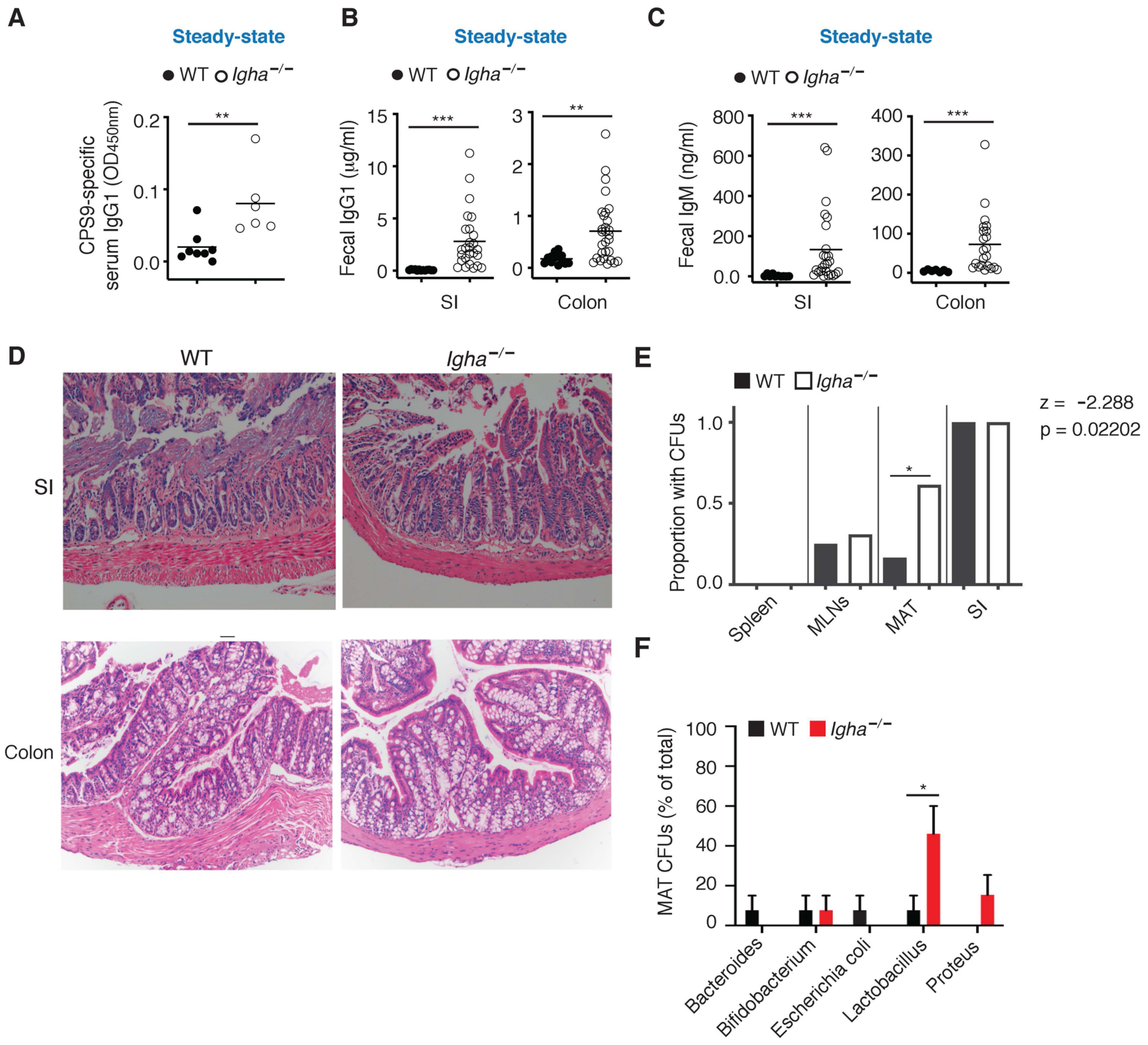


**Figure S4 (Related to Figure 4). IgA Restrains Systemic IgG Responses to Translocated Gut Antigens**

(A) ELISA of serum IgG1 to pneumococcal CPS9 from 8 WT or 6 *Igha*$^{-/-}$ mice at steady state.

(B) ELISA of intraluminal free IgG1 from the SI or colon feces of 12-13 WT or 26-29 *Igha*$^{-/-}$ mice at steady state.

(C) ELISA of free IgM from the SI or colon feces of 7-10 WT or 22-25 *Igha*$^{-/-}$ mice at steady state.

(D) Hematoxylin and eosin staining of tissue sections from the SI and colon of representative WT or *Igha*$^{-/-}$ mice at steady state (pathological score 0).

(E) Overall percentage of agar plates from each organ homogenate that resulted in the identification of CFUs after incubation in an anaerobic chamber. Results for spleen, MLN, and MAT are from 12-13 WT and 13 *Igha*$^{-/-}$ mice, while positive controls from the small intestine are from 7 WT and 10 *Igha*$^{-/-}$ mice. Based on the same data as in Figure 4F.

(F) Summary of CFUs of anaerobic bacteria identified in homogenates of MAT from 12 WT or 13 *Igha*$^{-/-}$ mice at steady state. The frequency (%) of colony-positive agar plates containing each taxonomically identified genus per experiment is shown.

Data derive from one experiment (A), summarize 4-5 experiments (B, C, E, F), or are representative of 4 WT and 4 *Igha*$^{-/-}$ mice (D). Data are presented with mean (A-C) or mean ± s.e.m. (F); significance was determined using two-tailed unpaired Mann-Whitney test (A, B, C, F) or a two-tailed two proportions z-test (E). *$p < 0.05$, **$p < 0.01$, ***$p < 0.001$.

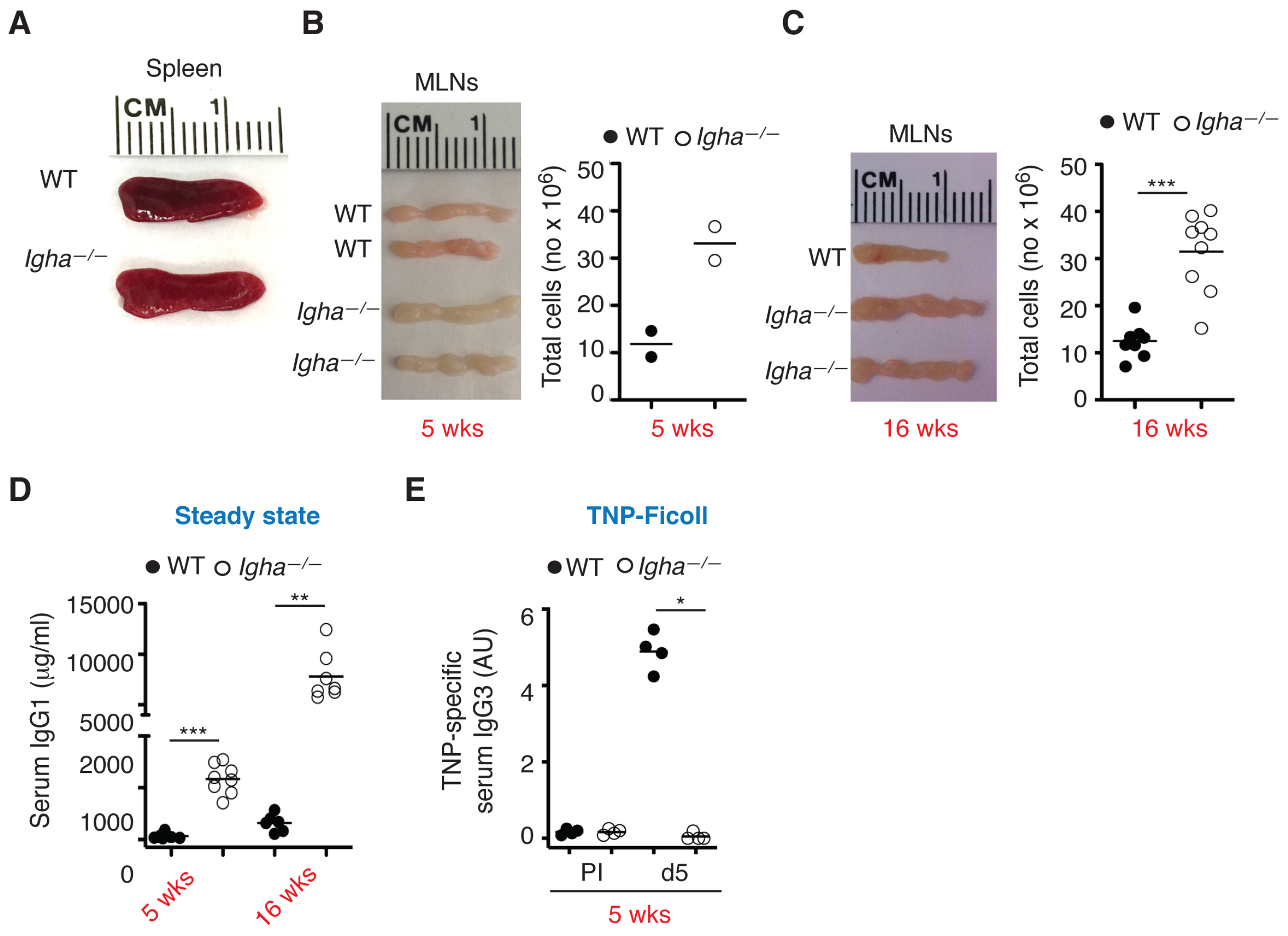


**Figure S5 (Related to Figure 5). IgA Starts Limiting Systemic IgG1 Production to Gut Microbial Antigens at a Very Early Age**

(A) Images of spleen from representative adult WT or *Igha*$^{-/-}$ mice at steady state.

(B) Left: images of mesenteric lymph nodes (MLNs) from two WT or *Igha*$^{-/-}$ mice aged 5 weeks at steady state. Right: absolute numbers of cells from MLNs of the same mice.

(C) Left: images of MLNs from a representative WT mouse or two *Igha*$^{-/-}$ mice aged 16 weeks at steady state. Right: absolute numbers of cells from MLNs of 8 WT or 9 *Igha*$^{-/-}$ mice aged 16 weeks.

(D) Total serum IgG1 from 6-7 WT or 7-8 *Igha*$^{-/-}$ mice aged 5 or 14 weeks at steady state.

(E) ELISA of serum IgG3 to TNP from 4 WT or *Igha*$^{-/-}$ mice aged 5 weeks prior to immunization (PI) and 5 days following i.p. immunization with TNP-Ficoll.

Data are from one representative experiment of 2-3 (A, C left), 2-3 experiments (C right, D right), or one individual experiment (B, E). Data are presented with mean; a two-tailed unpaired Student's t-test was performed when data were determined to follow a Gaussian distribution (D), otherwise a Mann-Whitney test was used to determine significance (C, E). *$p < 0.05$, **$p < 0.01$, ***$p < 0.001$.

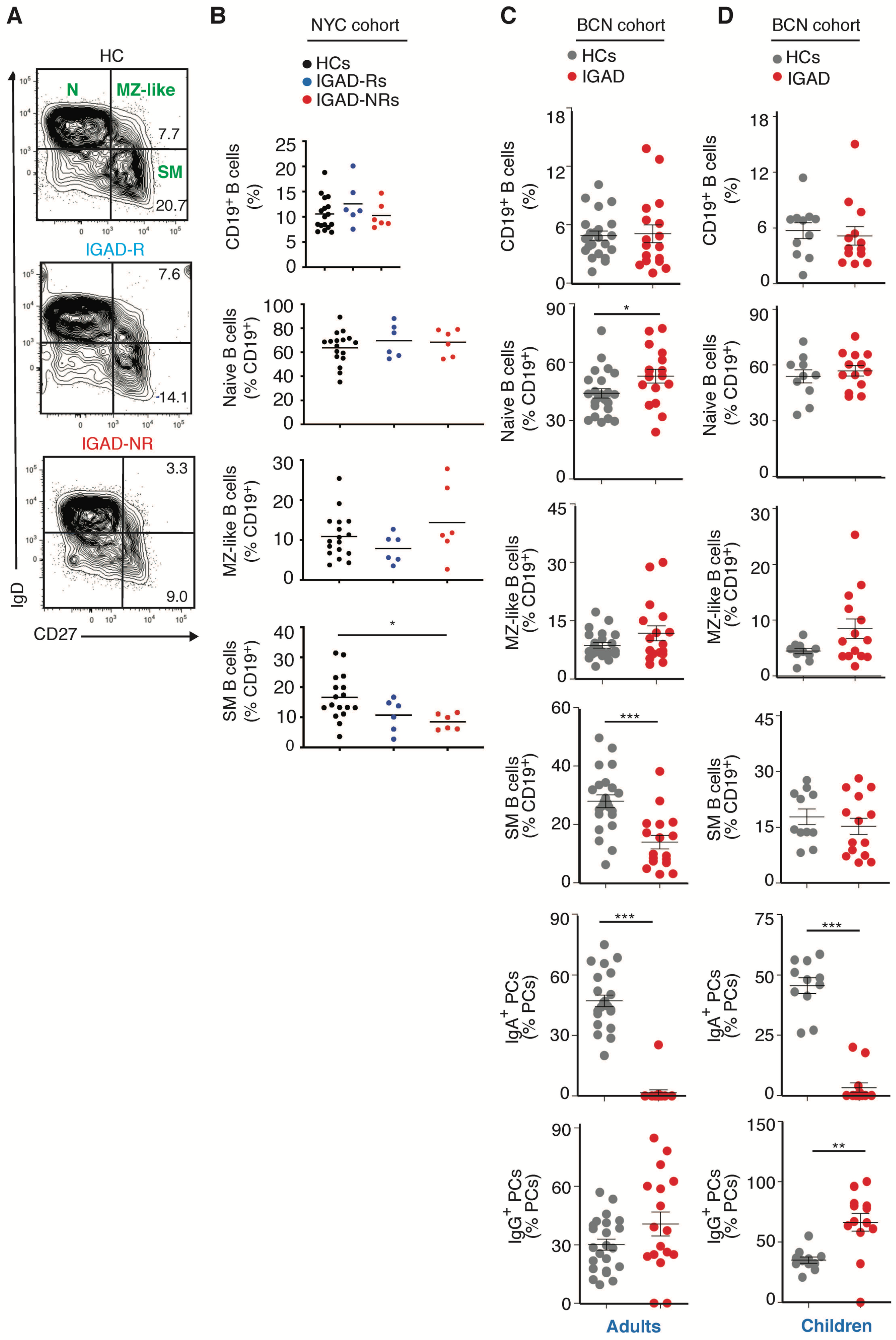


**Figure S6 (Related to Figure 6). IgA Sustains GC-Derived Switched Memory B Cells in Humans**

(A) Flow cytometry analysis of IgD and CD27 on circulating naïve (N) $IgD^{+}CD27^{-}$, MZ-like $IgD^{+}CD27^{+}$ and switched memory (SM) $IgD^{-}CD27^{+}$ B cells in a representative adult healthy control (HC) donor and two representative adult IGAD patients from the NYC cohort, including adult IGAD-Rs and IGAD-NRs defined by the presence or absence of an adequate IgG response to vaccination with Pneumovax23. Numbers indicate frequency (% of $CD19^{+}$ cells).

(B) Frequency of circulating total $CD19^{+}$ B cells (% of live), naïve, MZ-like and SM B cell subsets (% of $CD19^{+}$) from adult donors of a NYC cohort comprised of 17 HCs, 6 IGAD-Rs and 6 IGAD-NRs.

(C) Frequency of circulating $CD19^{+}$ total B cells (% of live), $IgM^{+}IgD^{+}CD27^{-}$ naïve B cells, $IgM^{hi}IgD^{int}CD27^{+}$ MZ-like B cells, $IgD^{-}IgM^{-}CD27^{+}$ SM B cells (% of $CD19^{+}$), $IgA^{+}$ PCs and $IgG^{+}$ ($IgM^{-}IgA^{-}$) PCs (% of $CD19^{+}CD27^{hi}CD38^{hi}$) from adult donors of a BCN cohort comprised of 23 HCs and 17 IGAD patients. All subsets were first gated on $CD19^{+}$non-$CD38^{++}CD27^{++}$ cells.

(D) Frequency of circulating $CD19^{+}$ total B cells (% of live), $IgM^{+}IgD^{+}CD27^{-}$ naïve B cells, $IgM^{hi}IgD^{int}CD27^{+}$ MZ-like B cells, $IgD^{-}IgM^{-}CD27^{+}$ SM B cells (% of $CD19^{+}$), $IgA^{+}$ PCs and $IgG^{+}$ PCs (% of $CD19^{+}CD27^{hi}CD38^{hi}$) from children of a BCN cohort comprised of 11 HCs and 14 IGAD patients, gated as in (C).

Data show a representative flow cytometry contour plot (A) or paired experiments wherein flow cytometry was performed on human peripheral blood mononuclear cells upon reception of IGAD blood samples paired with HC blood samples (B-D). Data are presented with mean or mean ± s.e.m.; significance was determined using two-tailed unpaired Mann-Whitney test or a Kruskal-Wallis test with Dunn's correction for multiple comparisons. *$p < 0.05$, ***$p < 0.001$.

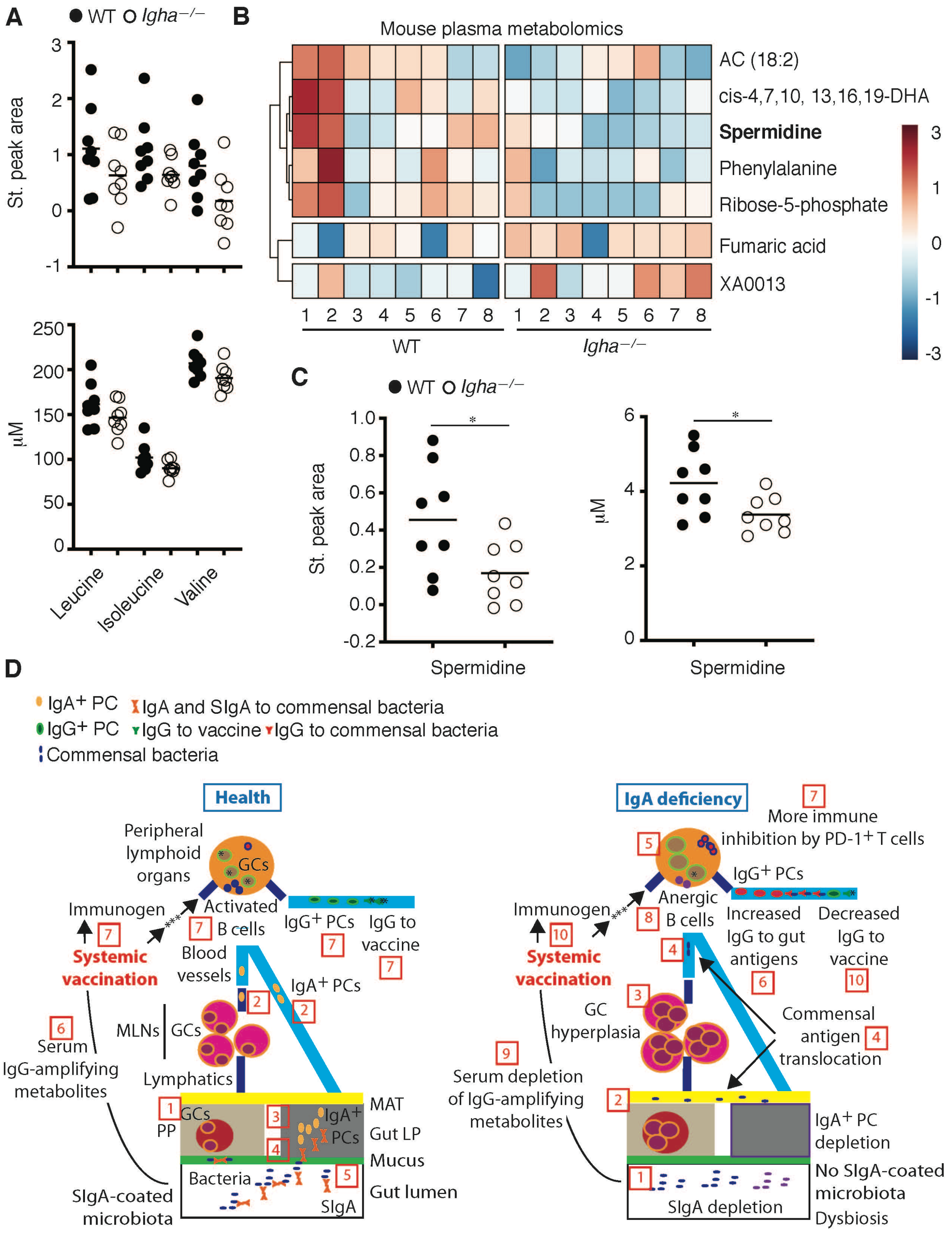


**Figure S7 (Related to Figure 7). IgA Regulates Systemic Metabolites.**

(A) Plasma concentration of leucine, isoleucine and valine from 8 WT or *Igha*$^{-/-}$ mice. Standardized (st.) peak area was compared to internal standard (top); quantitative estimation (μM) was performed by normalizing relative peak areas and comparing to standard curve (bottom).

(B) Heat map of plasma metabolomics from 8 WT or *Igha*$^{-/-}$ mice. Significantly different metabolites are shown and spermidine is highlighted in bold. AC, acylcarnitine; DHA, docosohexahenoic acid.

(C) Plasma concentration of spermidine from 8 WT or *Igha*$^{-/-}$ mice. St. peak area was compared to internal standard (left); quantitative estimation (μM) was performed by normalizing relative peak areas and comparing to standard curve (right).

(D) Proposed model depicting the impact of gut IgA on systemic IgG responses to pneumococcal vaccines. **LEFT**. In healthy individuals, (1) bacteria-reactive B cells from IgA inductive sites such as PP-associated GCs generate $IgA^+$ PCs, which (2) reach the general circulation via the lymphatic system to (3) home back to the intestinal LP. Here, $IgA^+$ PCs release dimeric IgA, which (4) crosses the epithelial barrier through the pIgR (not shown) to become SIgA. In the intestinal lumen (5), SIgA binds to the commensal microbiota to inhibit bacterial translocation into tissues (i.e., immune exclusion), retain beneficial bacteria into the mucus layer (i.e., immune inclusion), and control bacterial function, including immune properties and metabolism. While immune inclusion contributes to gut homeostasis, immune exclusion prevents uncontrolled B and T cell activation by abnormally translocated commensal antigens. Of note, IgA-coated gut bacteria sustain the peripheral supply of metabolites (6) such as BCAAs with IgG-amplifying function. In this positive immune environment, systemic inoculation of pneumococcal vaccines (7) generates IgG-releasing PCs that mount protective IgG responses to the vaccine. **RIGHT**. In IgA-deficient individuals, IgA (1) does not coat gut bacteria, which (2) causes abnormal microbial translocation into the MAT. The ensuing enhanced translocation of commensal antigens (3) triggers excessive activation of the intestinal immune system, including B and T cells from PPs and MLNs. As a result, follicles from both PPs and MLNs develop GC hyperplasia. Gut commensal antigens further (4) translocate into the circulation and reach peripheral lymphoid tissues, thereby (5) causing additional GC hyperplasia and hyperactivation of systemic T and B cells. While hyperactivated B cells (6) differentiate into plasma cells secreting IgG antibodies to gut commensal antigens, hyperactivated T cells (7) expand and express more PD-1, which delivers negative immune signals after engaging PD-L1 on B cells (not shown). This immune inhibition (8) progressively leads to B cell anergy and is compounded by depletion of gut microbiota-controlled metabolites such as BCAAs with IgG-amplifying functions (9). In this negative immune environment, systemic inoculation of pneumococcal vaccines (10) fails to induce sufficient IgG responses to TD or TI pneumococcal vaccines.
